# Predicting Primary Graft Dysfunction in Lung Transplantation: Machine Learning–Guided Biomarker Discovery

**DOI:** 10.1101/2024.05.24.595368

**Authors:** Dianna Nord, Jason Cory Brunson, Logan Langerude, Hassan Moussa, Blake Gill, Tiago Machuca, Mindaugas Rackauskas, Ashish Sharma, Christine Lin, Amir Emtiazjoo, Carl Atkinson

## Abstract

**BACKGROUND:** There is an urgent need to better understand the pathophysiology of primary graft dysfunction (PGD) so that point-of-care methods can be developed to predict those at risk. Here we utilize a multiplex multivariable approach to define cytokine, chemokines, and growth factors in patient-matched biospecimens from multiple biological sites to identify factors predictive of PGD.

**METHODS:** Biospecimens were collected from patients undergoing bilateral LTx from three distinct sites: donor lung perfusate, post-transplant bronchoalveolar lavage (BAL) fluid (2h), and plasma (2h and 24h). A 71-multiplex panel was performed on each biospecimen. Cross-validated logistic regression (LR) and random forest (RF) machine learning models were used to determine whether analytes in each site or from combination of sites, with or without clinical data, could discriminate between PGD grade 0 (*n* = 9) and 3 (*n* = 8).

**RESULTS:** Using optimal AUROC, BAL fluid at 2h was the most predictive of PGD (LR, 0.825; RF, 0.919), followed by multi–timepoint plasma (LR, 0.841; RF, 0.653), then perfusate (LR, 0.565; RF, 0.448). Combined clinical, BAL, and plasma data yielded strongest performance (LR, 1.000; RF, 1.000). Using a LASSO of the predictors obtained using LR, we selected IL-1RA, BCA-1, and Fractalkine, as most predictive of severe PGD.

**CONCLUSIONS:** BAL samples collected 2h post-transplant were the strongest predictors of severe PGD. Our machine learning approach not only identified novel cytokines not previously associated with PGD, but identified analytes that could be used as a point-of-care cytokine panel aimed at identifying those at risk for developing severe PGD.

## 1 Introduction

Lung transplantation is an acceptable therapy for end-stage lung disease. However, successful posttransplantation outcomes are significantly impacted by the development of Primary Graft Dysfunction (PGD). Severe PGD is associated with early mortality and a more rapid onset of chronic lung allograft dysfunction (CLAD) (Christie et al. 2010; Huang et al. 2008; Daud et al. 2007; Kreisel et al. 2011). As such, there is an urgent need to better understand PGD pathobiology and identify those at enhanced risk. The pathophysiology of PGD is complex and involves a multifactorial inflammatory process that likely starts in the donor lung as a consequence of donor injury and cold storage that is subsequently exacerbated by ischemia reperfusion injury (IRI).

Given the importance of inflammation in mediating PGD, a number of studies have sought to leverage the presence of pro-inflammatory cytokines as prognostic biomarker targets that can assist in stratifying disease severity and guide clinical management. Previous studies looking for informative biomarkers in PGD have largely done so through measuring the presence of soluble factors in recipient plasma. Multiple studies have demonstrated that serum elevation of pro-inflammatory cytokines such as IL-6, IL-8, IP-10, and MCP-1 are associated with severe PGD (Moreno et al. 2007; Chacon-Alberty et al. 2022; Mathur et al. 2006; Hoffman et al. 2009; Bharat et al. 2008; Shah et al. 2012; Bonneau et al. 2022; Allen et al. 2012). Furthermore, other studies have identified elevated IL-10 in PGD+ plasma post-transplant, suggesting a paradoxical balance between pro- and anti-inflammatory markers in the post-transplant lung that is not yet fully understood (Mathur et al. 2006; Allen et al. 2012). In addition to investigating inflammatory signals, a study utilizing samples from the Lung Transplant Outcomes Group (LTOG) cohort assessed markers specific to endothelial injury, epithelial injury, and the coagulation cascade. These analyses found increased plasma PAI-1, ICAM, sRAGE, and SP-D and decreased plasma Protein C correlated to PGD development, which were predictive of 90-day mortality (Christie et al. 2007, 2010; Shah et al. 2012). However, these markers were not found to correlate with PGD in subsequent studies, indicating a need to identify optimal common biomarkers (Chacon-Alberty et al. 2022).

Although recipient plasma provides a valuable window into the biological/inflammatory processes detected on a systemic level, a handful of studies have analyzed bronchoalveolar lavage (BAL) fluid as a more representative sample of the post-transplant lung microenvironment. Three independent studies, which analyzed BAL fluid collected 24 hours post-transplantation from PGD+ patients, demonstrated increased levels of IL-6, IL-8, IL-10, IL-13, Eotaxin, G-CSF, IFN-C, MIP-1α, TGFβ, and procollagen type I (PC) as compared to BAL from PGD-free patients (Frick et al. 2020; Verleden et al. 2018; DerHovanessian et al. 2016). These markers suggest interplay between pro- and anti-inflammatory cytokines, innate immune chemoattractants, and mediators of fibroproliferation in the lung microenvironment of PGD patients. In keeping with a focus on lung specific immune changes, recent ex vivo lung perfusion studies have assayed the perfusion solution pre-transplant and demonstrated that elevated IL-8, IL-6, and GRO-α levels correlate with poorer graft outcomes post transplantation (Machuca et al. 2015; Ghaidan et al. 2022).

Taken together, these studies have shown the potential value of measuring inflammatory and injury markers as predictive biomarkers. Most factors identified to date have been selected for investigation by association, in that they have been previously implicated in other acute lung injuries and subsequently investigated in lung transplantation. While these have guided our understanding of PGD, their hypothesis driven nature may limit the identification of specific mediators or predictors of PGD. Here, we utilized an expansive 71-analyte multiplex approach composed of a broad array of cytokine, chemokines, and growth factors in concert with matched donor perfusate, recipient plasma, and BAL fluid as a means to identify novel biomarkers and mechanistic factors associated with PGD. By examining multiple analytes across diverse samples, our study aimed to mitigate selection bias and uncover stronger correlates to PGD development. We further hypothesized that a machine learning approach may serve as a valuable tool that complements classical statistics for biomarker discovery. Predicting PGD onset using machine learning has been previously explored using donor and recipient clinical characteristics and/or analysis of exhaled compounds during spirometry (Stefanuto et al. 2022; Michelson et al. 2023), but has yet to be utilized in multiplex analyte discovery studies. Here, we used a machine learning approach to predict PGD from multi-site multiplex data and identify key predictors as candidate biomarkers. We found that cytokine expression in BAL fluid samples collected 2hr post-transplant best predicted PGD development. Prediction was enhanced by using clinical indices including donor and recipient characteristics. A model reduction process obtained 3 BAL fluid predictors of PGD development: BCA-1 (CXCL13), Fractalkine, and IL-1Ra, factors that have not been extensively studied in the context of lung transplantation. Taken together, these results provide a methodological foundation for further development of predictive modeling that could result in clinically relevant point-of-care testing and provide novel targets for mechanistic studies of PGD.

## 2 Methods

### 2.1 Inclusion/Exclusion Criteria

Samples were acquired from the University of Florida Transplant Tissue Bank from patients undergoing a first-time bilateral lung transplant at The University of Florida between May 2021 and May 2022. Nineteen patients were selected based on those that had a complete sample set and that had the same PGD grade at all measured timepoints, either 0 (*n* = 10) or 3 (*n* = 9). PGD was graded as previously described using the International Heart and Lung Society (ISHLT) Criteria (Snell et al. 2017). Donor and recipient demographics for each group are detailed in Table 1. This study was approved by the University of Florida Institutional Review Board IRB#202200536.

**Table 1.** Baseline donor and recipient characteristics. Donor and Recipient characteristics by primary graft dysfunction status. Differences between groups in categorial variables were analyzed by Fisher’s Exact Comparison and differences in numerical variables were analyzed by unpaired Student’s *t*-test, with *p <* 0.05 considered significant. ILD: Interstitial Lung Disease, IPF: Idiopathic Pulmonary Fibrosis, COPD: Chronic Obstructive Pulmonary Disease, LAS: Lung Allocation Score, ECMO: Extracorporeal Membrane Oxygenation.

| **Donor Characteristics** | **PGD- (*n* = 9)** | **PGD+ (*n* = 8)** | **p-value** |
| --- | --- | --- | --- |
| **Median Donor Age** | 37 (16, 60) | 37 (18, 60) | .974 |
| **Donor Sex** |  |  |  |
| Male | 89% (8) | 25% (2) | **.015** |
| Female | 11% (1) | 75% (6) |  |
| **Donor Race/Ethnicity** |  |  |  |
| White | 44% (4) | 50% (4) | – |
| Hispanic/Latino | 33% (3) | 38% (3) |  |
| Black or African American | 22% (2) | – |  |
| American Indian | – | 13% (1) |  |
| **Static Storage** |  |  |  |
| Standard 4°C | 100% (9) | 50% (4) | **.029** |
| 10°C | – | 50% (4) |  |
| **Smoking History** |  |  |  |
| Yes | 56% (5) | 38% (3) | .637 |
| No | 44% (4) | 63% (5) |  |
| **Heavy Alcohol Use History** |  |  |  |
| Yes | 22% (2) | 13% (1) | .999 |
| No | 78% (7) | 88% (7) |  |
| **Recipient Characteristics** | **PGD- (*n* = 9)** | **PGD+ (*n* = 8)** | **p-value** |
| **Median Recipient Age** | 61 (48, 74) | 59 (35, 73) | .418 |
| **Mean Recipient BMI at Listing** | 24.9 | 26.9 | .256 |
| **Recipient Sex** |  |  |  |
| Male | 78% (7) | 13% (1) | **.015** |
| Female | 22% (2) | 88% (7) |  |
| **UNOS Lung Disease Diagnosis Group** |  |  |  |
| Group A: Obstructive Lung Disease | 3 | 0 | – |
| Group B: Pulmonary Vascular Disease | 1 | 0 |  |
| Group C: Cystic Fibrosis | 0 | 0 |  |
| Group D: Restrictive Lung Disease | 5 | 8 |  |
| **Mean LAS Score** | 47.9 (33.1, 92.4) | 57.1 (36.5, 93.3) | .361 |
| **% ECMO Pre-Tx** | 10% (1) | 22% (2) | .577 |
| **% ECMO Post-Tx** | 10% (1) | 67% (5) | **.050** |
| **% Sensitization Protocol** | – (0) | 50% (4) | **.030** |

### 2.2 Sample Collection

Herein we analyzed three distinct sample types: (1) Plasma, (2) Bronchoalveolar lavage fluid (BAL), and (3) Donor lung organ perfusate solution. Plasma was processed from whole blood samples collected from lung transplant recipients within 12hr prior to transplant (pre), and at 2hr, 24hr, 72hr, 1wk, and 2wk post-transplant using SepMate tubes and Lymphoprep Solution (Stem Cell Technologies, USA) according to manufacturer’s recommended protocol. BAL fluid was obtained following reperfusion, at a timepoint matched to the 2hr blood sample, “T0”. Bronchoscopy and BAL fluid collection was performed in accordance with the ISHLT recommendations (Martinu et al. 2020). BAL fluid was processed by centrifugation, and supernatants were collected. Donor lung Perfadex® (Xvivo, Sweden) perfusate solution was obtained by collecting the initial volume of 2–3 mL of preservation solution passed out of the lung vasculature at the time of reperfusion and processed by centrifugation, and supernatants collected. All processed samples were stored at –80°C until time of analysis.

### 2.3 Multiplex assays

Analysis of patient plasma, BAL fluid, and perfusate was performed using commercially available Millipore Sigma MILLIPLEX® base kits as described below, on Luminex 100/200™ instruments (Luminex/DiaSorin, Saluggia, Italy), with Bio-Plex Manager™ (BPM) software (BioRad, Hercules, CA) by Eve Technologies (Calgary, Canada). Eve Technologies’ Cytokine, Chemokine, Growth Factor 71-Plex Clinical RUO Test (HD71-CLIN) was used to detect plasma, BAL fluid, and perfusate levels of 71 analytes using two separate panels as per the manufacturer’s instructions for use: (1) the Human Cytokine 48-Plex Discovery Assay® (HD48; Millipore MILLIPLEX® Human Cytokine/Chemokine/Growth Factor Panel Immunology Multiplex Assay, Cat. #HCYTA-60K), and (2) the Human Cytokine 23-Plex Discovery Assay® (HD23; Millipore MILLIPLEX® Human Cytokine/Chemokine Magnetic Bead Panel II Immunology Multiplex Assay, Cat. #HCYP2MAB-62K) (for the full analyte list, see Supplemental Table 1). Each specimen on each panel was run in singlet on the same analyzers by Eve Technologies, which is certified to perform high complexity laboratory testing under the Clinical Laboratory Improvement Amendments (CLIA). Since a relative absence (or low levels) of any particular analyte could be meaningful with respect to patterns of cytokine expression, any sample with a fluorescence value below the range of its standard curve (for which concentration values are unable to be interpolated) was included in the analysis by assigning a concentration value corresponding to the minimum sample value for the same analyte. Assay range values for each analyte are listed in Supplemental Table 1.

### 2.4 Software

Computations were performed in R (R Core Team 2021), using Tidyverse analysis workflows (Wickham et al. 2019) and Tidymodels modeling workflows (Kuhn and Wickham 2020). Several packages were used for specific analyses (Wickham and Bryan 2023; Tierney, Cook, et al. 2023; Tierney, Hughes, et al. 2023; Kolde 2019; Wei et al. 2021; Brunson and Paul 2022; Dahl et al. 2019; Greenwell and Boehmke 2023; Pedersen 2024a, 2024b; Galili et al. 2023; Kaplan and Schlegel 2023).

### 2.5 Preprocessing

Donor and recipient characteristics are displayed in Table 1. Smoking and alcohol history were collapsed to logical indicators of past use. We applied the offset log-transformation *x↦* log(*x* + 1) to observed concentrations before all analyses, and we refer to these values as “log-concentrations”. Research IDs were uniquely coded to capital letters.

### 2.6 Protein concentrations and pairwise associations

We visualized protein co-concentration patterns within each medium using heatmaps annotated by case and site dendrograms (Kolde 2019) and using correlation plots (Wei et al. 2021).

We used Kruskal–Wallis (KW) tests to identify proteins whose concentrations in plasma changed over time and included these proteins in a principal components analysis (PCA). For clarity, we plotted segments between consecutive time points of each case, and we overlaid segments between the centroids at each timepoint to produce an average trajectory.

### 2.7 Discrimination of PGD

We used KW (perfusate and BAL fluid) and multivariate Cramér (Baringhaus and Franz 2004) (plasma, all time points) tests to identify proteins that discriminated between the PGD– and PGD+ subgroups. We visually compared log-concentrations in each medium between the subgroups using jitter (perfusate and BAL fluid) and line (plasma) plots. We also generated a biplot of trajectories in plasma with PGD-stratified average trajectories overlaid.

### 2.8 Prediction of PGD

We used logistic regression, random forest, and nearest neighbors models in a machine learning (ML) workflow to determine whether several sets of predictors could discriminate between PGD– and PGD+ cases. To better reflect data availability in practice conditions, only concentrations calculated by Eve were used for predictive analysis; no missing values were imputed, so analytes with several missing values were excluded. The data pre-processing recipe is detailed in the Methodological Supplement.

The model families were optimized over hyperparameter grids. For each model specification, we calculated the area under the receiver operating characteristic curve for leave-one-out cross-validation (LOOCV), stratified by outcome so that each test set comprised one PGD– and one PGD+ case. We then repeated this procedure 12 times to ensure diversity in the test sets.

We assessed the predictive value of every set of data from 4 sites: perfusate, BAL fluid, first three time points of plasma, and clinical criteria (donor, recipient, and perioperative). We ranked the model specifications by mean AUROC across the 12 LOOCVs. We summarized the means and standard errors of the AUROCs as line–range plots and tabulated the optimal and mean near-optimal performance of each model family using data from each combination of sites.

Next, optimized model specifications were fitted to complete data from each combination of sites. We used absolute coefficient estimates (LR) and mean decrease in accuracy (RF) to rank predictors in each model family and data set. To test the feasibility of a point-of-care test, we included the top predictors from both families in a LR LASSO using data from the site that yielded the best predictions. Informed by the results, we selected a subet of proteins as a candidate biomarker panel.

### 2.9 Robustness checks

We performed three robustness checks on our results by re-running the full analysis after each of three alternative pre-processing choices: (1) for exploratory analysis, not imputing values when observed concentrations were not calculated, and for predictive analysis, using imputed values; (2) using fluorescence intensities (FIs), for which very few values were missing, in place of observed analyte concentrations; and (3) including two additional cases, one in each PGD group, for which pilot multiplex assays were performed in a separate batch.

## 3 Results

### 3.1 Sample characteristics and group differences

Seventeen patients were included in the complete analysis that had a complete sample set and that had PGD grade of 0 (n=9) or 3 (n=8) at all measured timepoints. In keeping with published clinical risk factors, patients in the PGD+ group were predominantly female and pre-sensitized. Interestingly, lungs stored at 10C were more likely to be PGD+ (Table 1).

### 3.2 Plasma protein concentrations and pairwise comparisons

We observed relatively consistent protein clustering based on concentrations of analytes in plasma over the first 3 time points (pre-Tx, T0, and 24h post-transplantation) (Figure 1A). We spotlight 4 protein clusters based on co-concentrations in plasma over the first 3 time points (pre-LTx, T0, and 24h) (Figure 1A). First, a high-expression cluster comprises PDGF-AB/BB, G-CSF, RANTES, MIG/CXCL9, SDF-1α+β, IL-27, MIP-1δ, sCD40L, and IL-17E/IL-25. The next bifurcation leads to two large clusters, one of which comprises the lowest-concentrated proteins: MCP-4, FLT-3L, IL-5, SCP, Eotaxin-3, I-309, IL-21, LIF, IL-3, TSLP, TNFβ, IL-33, IL-7, IL-2, IFNγ, IL-4, IL-17A, TGFα, IL-17F, IL-1β, IL-12p70, IL-15, IL-9, and IL-28A. Finally, the remaining branch bifurcates into two moderately concentrated subgroups. The less-concentrated of these comprises IL-1RA, IL-6, IL-10, IL-23, TPO, IL-12p40, M-CSF, IL-13, IL-22, IFN-α2, MCP-3, MIP-1α, TARC, EGF, VEGF-A, GROα, IL-8, TRAIL, Eotaxin, and MCP-2. The more-concentrated comprises GM-CSF, IL-1α, TNFα, FGF-2, Fractalkine, IP-10, MIP-1β, BCA-1, CTACK, IL-18, MCP-1, 6CKine, MDC,Eotaxin-2, IL-16, PDGF-AA, and ENA-78. One outlier case is evident (G) with higher levels of proteins in low-concentration clusters. With only 6 other exceptions, the panels cluster neatly by time point.

**Figure 1.**
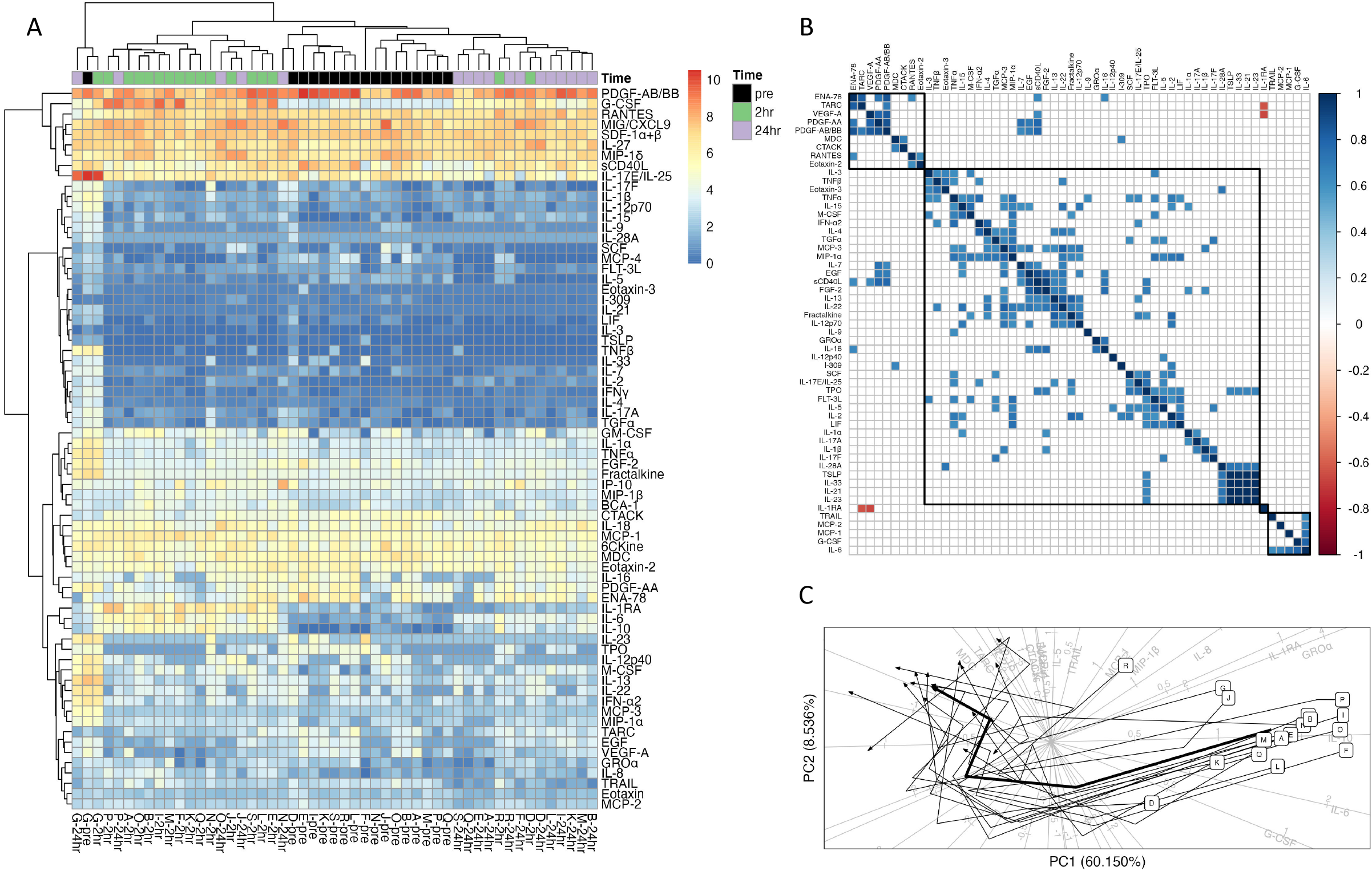
Log-concentrations of 71 proteins in plasmaat pre-Tx, T0, and 24h post-transplantation. Protein levels and their changes during the peri-operative period were similar across all but one subject. (A) Clustered heatmap for all subjects at pre-Tx, T0, and 24h time points, using complete-linkage clustering on cases and proteins. Blue (red) indicates lower (higher) levels. (B) Correlation matrix of soluble factors measured in plasma from all subjects at T0. Only correlations with *p <* 0.01 in Spearman tests are filled. Proteins are ordered by complete-linkage clustering in order to juxtapose more correlated proteins. Note that complete-linkage clustering is heuristic, so the order may differ from that of (A). (C) PCA biplot of time-discriminating proteins at all subject–time points. Values at pre-Tx are labeled by unique keys. Segments connect adjacent time points for each subject. Segments are grouped by PGD status. Thickened segments connect time point centroids, stratified by PGD status.

We selected a distance threshold based on visual inspection of dendrograms to identify clusters in the correlation heatmap at T0 (Figure 1B). The resulting clusters were one singleton, IL-1RA, the only protein to be detectably negatively correlated with other proteins (TARC and VEGF-A); a small cluster containing its negative correlates as well as ENA-78, PDGF-AA, PDGF-AB/BB, MDC, CTACK, RANTES, and Eotaxin-2; another small cluster comprising TRAIL, MCP-2, MCP-1, G-CSF, and IL-6; and the remaining proteins in a single large cluster.

The PCA revealed two distinct dimensions of variation in plasma protein concentrations over time, which captured nearly 70% of the total variance (Figure 1C). IL-1RA, GROα, IL-10, IL-6, and G-CSF loaded more strongly onto PC1, all positively. As patients proceeded from pre-Tx to 24h, PC1 decreased, then remained at lower levels through 2wk. MDC, TARC, IL-3, MIP-1δ, CTACK, Eotaxin, IL-2, IL-5, and TRAIL loaded more strongly onto PC2, MDC and TARC negatively. PC2 decreased from pre-Tx to T0, then increased gradually through 2wk. MCP-1, MIP-1β, and IL-8 were more collinear with each other than with either of these groups; on average, they loaded similarly onto both PCs. These patterns are reflected in the average trajectory.

### 3.3 Plasma protein discrimination of PGD

We next analyzed whether protein changes in the in plasma differed by PGD status. Analysis identified two to four clusters at T0, which broadly agree and exhibit similar concentration levels over the PGD– and PGD+ subgroups (Figure 2A). Similarly, mean trajectories in the PCA did not differ systematically between the PGD– and PGD+ groups (Figure 2B). We did find evidence of systematically different concentrations of certain proteins at different time points (Table 2), notably BCA-1 at 1wk and 6CKine at T0. Multivariate Cramér tests detected only two proteins with different concentration trajectories over 6 time points (Table 3; Figure 2C): BCA-1 and IL-22 (p < 0.05), both with higher concentrations in the PGD+ group, but similar overall trajectories.

**Table 2.** Results of Kolmogorov–Smirnov tests of differences between PGD groups in protein concentrations in plasma at single time points. Analytes are listed by sample time point first and in order of statistical evidence, up to *p <* 0.05.

| Time | Protein | $n_0$ | $n_3$ | Statistic | p-value |
| --- | --- | --- | --- | --- | --- |
| pre | BCA-1 | 9 | 8 | 0.667 | 0.02024 |
| 2hr | 6CKine | 9 | 8 | 0.778 | 0.00255 |
| 2hr | IL-22 | 9 | 8 | 0.778 | 0.00428 |
| 2hr | IL-16 | 9 | 8 | 0.750 | 0.01119 |
| 2hr | MCP-4 | 9 | 8 | 0.542 | 0.04977 |
| 24hr | 6CKine | 9 | 8 | 0.778 | 0.00428 |
| 24hr | IL-6 | 9 | 8 | 0.764 | 0.00831 |
| 24hr | IL-22 | 9 | 8 | 0.667 | 0.02024 |
| 24hr | MCP-1 | 9 | 8 | 0.667 | 0.02024 |
| 24hr | IL-15 | 9 | 8 | 0.639 | 0.04689 |
| 24hr | BCA-1 | 9 | 8 | 0.639 | 0.04689 |
| 72hr | I-309 | 9 | 8 | 0.764 | 0.00831 |
| 72hr | PDGF-AA | 9 | 8 | 0.667 | 0.02024 |
| 72hr | IL-12p40 | 9 | 8 | 0.556 | 0.02941 |
| 72hr | BCA-1 | 9 | 8 | 0.653 | 0.03357 |
| 72hr | IL-3 | 9 | 8 | 0.556 | 0.04689 |
| 72hr | IL-6 | 9 | 8 | 0.639 | 0.04689 |
| 1wk | BCA-1 | 9 | 8 | 0.875 | 0.00140 |
| 1wk | TRAIL | 9 | 8 | 0.750 | 0.01119 |
| 1wk | PDGF-AB/BB | 9 | 8 | 0.653 | 0.03357 |
| 1wk | MIP-1δ | 9 | 8 | 0.653 | 0.03357 |

**Table 3.** Results of multivariate Crámer tests of differences between PGD groups in protein concentrations across all time points. Analytes are listed in order of statistical evidence, up to *p <* 0.1.

| Protein | $n_0$ | $n_3$ | Statistic | p-value |
| --- | --- | --- | --- | --- |
| BCA-1 | 42 | 42 | 5.007 | 0.0050 |
| IL-22 | 42 | 42 | 4.876 | 0.0240 |
| IL-15 | 42 | 42 | 1.646 | 0.0609 |
| 6CKine | 42 | 42 | 1.892 | 0.0829 |
| Eotaxin | 42 | 42 | 0.850 | 0.0899 |

**Figure 2.**
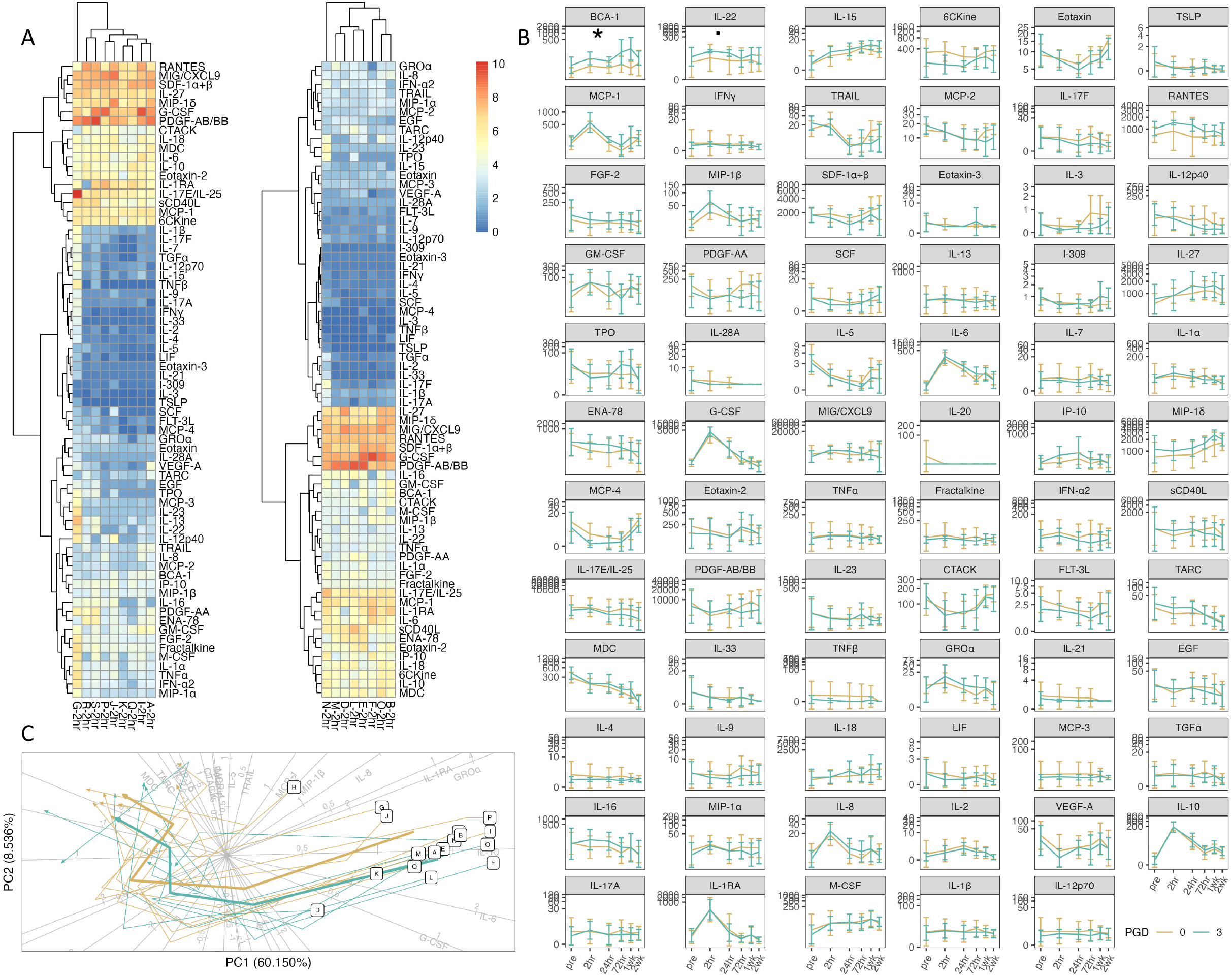
Differences in protein log-concentrations in plasma by PGD grade. Time series of protein levels did not detectably differ between PGD– and PGD+ subjects. (A) Clustered heatmaps for PGD– and PGD+ subjects at T0, using complete-linkage clustering on cases and proteins. (B) PCA biplot of time-discriminating proteins at all subject–time points, group-averaged by PGD grade (compare to Figure 1C). (C) Mean concentration and 1–standard deviation intervals across all time points, grouped by PGD grade. The panels are in order of increasing multivariate Cramér test p-value and marked: \*\*\**p <* 0.0001; \*\**p <* 0.001; \**p <* 0.01; _·_*p <* 0.05. See the Methodological Supplement for a table of p-values. Note that tests detected differences in BCA-1 and IL-22 between PGD– and PGD+, though this number (2) is expected by chance. Irrespective of PGD grade, cytokines and chemokines peaked 24h post-transplantation in response to ischemia reperfusion injury.

A subset of proteins (MCP-1, MIP-1β, IL-6, G-CSF, GROα, IL-8, IL-10, IL-1RA) spiked at T0 and then regressed toward their levels pre-LTx, dynamics that were not dissimilar between PGD– and PGD+ patients. These loaded strongly onto PC1 (Figure1C). For most other proteins, inter-case variation, as indicated by the intervals, was high relative to variation over time, though this does not take into account that variation over time was correlated across cases.

### 3.4 Perfusate protein discrimination of PGD

The donor organ is subjected to a host of injurious events prior to implantation. Brain death, organ procurement and cold storage can all result in injuries that shape early graft injuries post transplantation. A recent study showed that, despite the physiological basis that cold storage reduces cellular functions, analysis of preservation solution taken from donor hearts pre-transplant contained a myriad of cytokines (Ledwoch et al. 2022). Based on these findings, and the absence of previous such studies in lung transplantation, we utilized the same protein panel we used for plasma analysis on the lung storage solutions collected at the time of transplantation.

Two proteins were consistently highly expressed in perfusate: FGF-2 and MIP-1δ. These clustered together with other higher-concentration proteins, while lower-concentration proteins formed a second, larger cluster (Figure 3A). The cases did not clearly cluster along perfusate concentrations; the most distinctive patterns arose from three outliers (S, P, I). Clustering that was observed did not correlate with PGD grade. A direct comparison of protein levels in all cases (Figure 3B) suggests that organs with higher analyte levels were associated with PGD+, but we found no evidence that any single protein discriminated between PGD– and PGD+ cases (Table 4). Of the 66 proteins with sufficient data to compare, only two KW tests resulted in *p <* 0.1 (IL-17A and IL-1RA), a number expected by chance. The PCA suggests that variation in overall cytokine profiles is not associated with PGD group (Figure 3C).

**Table 4.**
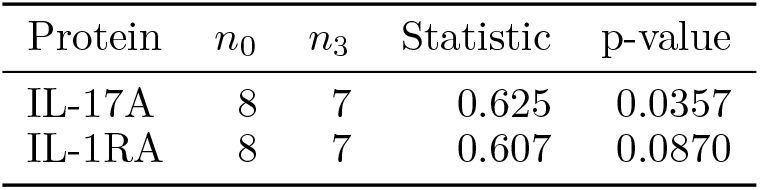
Results of Kolmogorov–Smirnov tests of differences between PGD groups in protein concentrations in perfusate. Analytes are listed in order of statistical evidence, up to *p <* 0.1.

**Figure 3.**
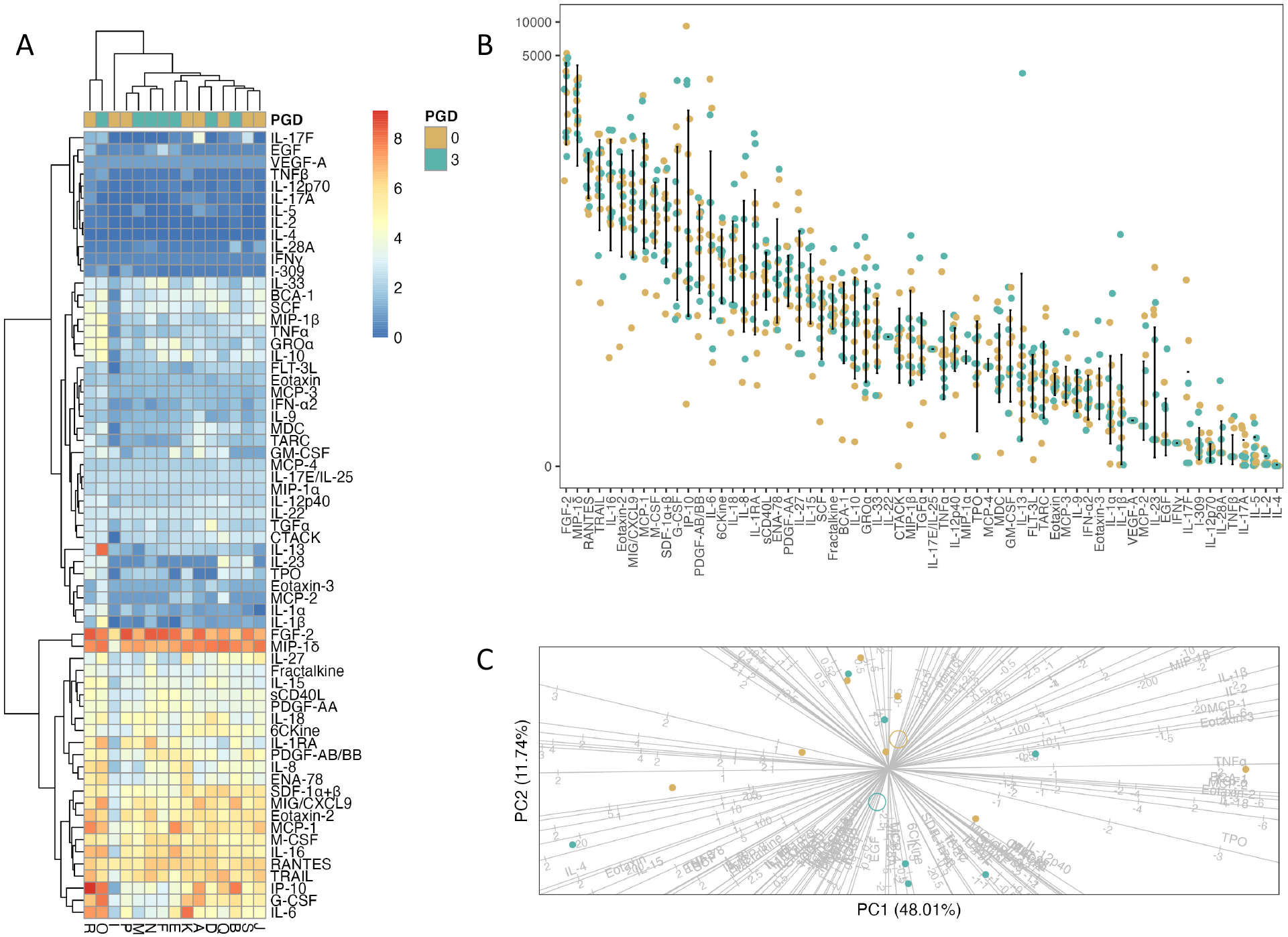
Differences in protein log-concentrations in perfusate by PGD grade. Protein levels did not detectably differ between PGD– and PGD+ subjects. (A) Clustered heatmap for all subjects at T0, using complete-linkage clustering on cases and proteins, annotated by PGD grade. (B) Jitter plots, grouped by PGD grade. Proteins are in order of decreasing median concentration. (C) PCA biplot on all available protein concentrations. PGD group averages are indicated by open circles.

### 3.5 BAL protein discrimination of PGD

The cases do not cluster more clearly in BAL fluid than in perfusate, nor do the clusters that are present appear to correspond to PGD grade (Figure 4A). As with perfusate, higher analyte concentrations generally seem to correspond to PGD+ (Figure 4B); in contrast to perfusate, this pattern was statistically evidenced for a small number of proteins (Table 5): IL-1RA (*p <* 0.01), Fractalkine, BCA-1, Eotaxin, IL-12p40, IL-6, IP-10, and M-CSF (*p <* 0.05). The PCA on all 70 proteins showed a clear separation between PGD– and PGD+ cases (Figure 4C), indicating that much of the variation in cytokine profiles is associated with severity of graft dysfunction.

**Table 5.**
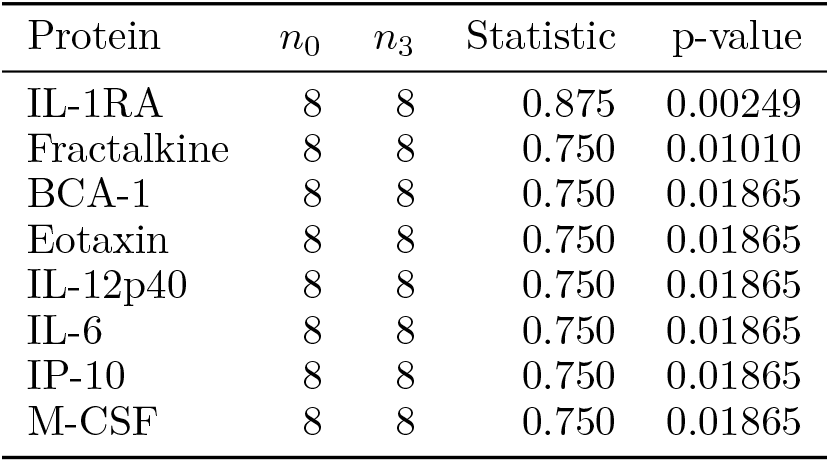
Results of Kolmogorov–Smirnov tests of differences between PGD groups in protein concentrations in BAL fluid. Analytes are listed in order of statistical evidence, up to *p <* 0.05.

| Protein | $n_0$ | $n_3$ | Statistic | p-value |
| --- | --- | --- | --- | --- |
| IL-1RA | 8 | 8 | 0.875 | 0.00249 |
| Fractalkine | 8 | 8 | 0.750 | 0.01010 |
| BCA-1 | 8 | 8 | 0.750 | 0.01865 |
| Eotaxin | 8 | 8 | 0.750 | 0.01865 |
| IL-12p40 | 8 | 8 | 0.750 | 0.01865 |
| IL-6 | 8 | 8 | 0.750 | 0.01865 |
| IP-10 | 8 | 8 | 0.750 | 0.01865 |
| M-CSF | 8 | 8 | 0.750 | 0.01865 |

**Figure 4.**
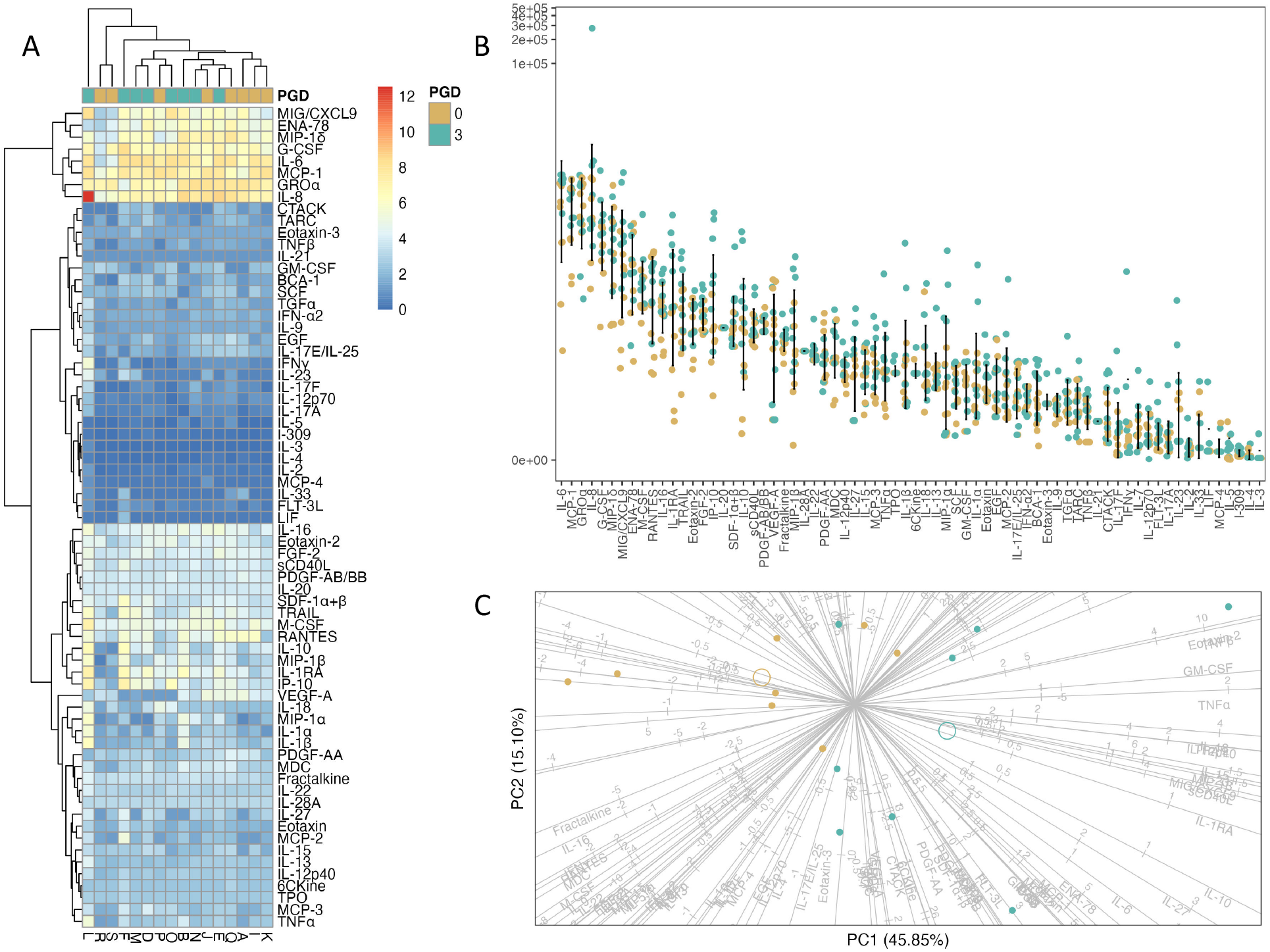
Differences in protein log-concentrations in BAL fluid by PGD grade; compare to Figure 3. Differences in the levels of several proteins were detected between PGD– and PGD+ subjects. (A) Clustered heatmap for all subjects at T0, using complete-linkage clustering on cases and proteins, annotated by PGD grade. (B) Jitter plots, grouped by PGD grade. Proteins are in order of decreasing median concentration. (C) PCA biplot on all available protein concentrations. PGD group averages are indicated by open circles.

### 3.6 Predictive modelling of PGD

We found weak statistical evidence for associations in specimens between analyte concentrations and PGD status, with the exception of BAL fluid. These results led us to consider predictive modeling via machine learning as an alternative approach.

Tables 6 and 7 compare model performance across each predictor set. Table 6 presents the highest AUROC obtained using each combination of data and each model family. To assess the sensitivity of these results to parameter tuning and moderate expectations for out-of-box performance, AUROCs in Table 7 are averaged from all models within 1 step on the hyperparameter grid from an optimal parameter setting (including optimal).

**Table 6.** Highest predictive performance obtained across parameter grids within each model family and each combination of predictors. Performance is measured by the area under the receiver operating characteristic curve.

| Perfusate | BAL | Plasma | $n_0, n_3$ | Multiplex | | | Multiplex and Clinical | | | |
| --- | --- | --- | --- | --- | --- | --- | --- | --- | --- | --- |
| | | | | LR | NN | RF | $n_0, n_3$ | LR | NN | RF |
|  |  |  |  |  |  |  | 10, 9 | 0.944 | 0.852 | 0.938 |
| × |  |  | 8, 7 | 0.565 | 0.514 | 0.448 | 8, 7 | 0.983 | 0.781 | 0.846 |
|  | × |  | 8, 8 | 0.825 | 0.681 | 0.919 | 8, 8 | 0.979 | 0.979 | 0.917 |
|  |  | × | 9, 8 | 0.841 | 0.441 | 0.653 | 9, 8 | 1.000 | 0.826 | 1.000 |
| × | × |  | 8, 7 | 0.812 | 0.710 | 0.889 | 8, 7 | 1.000 | 0.785 | 0.778 |
| × |  | × | 8, 7 | 0.500 | 0.250 | 0.525 | 8, 7 | 0.965 | 0.667 | 0.944 |
|  | × | × | 8, 8 | 0.833 | 0.417 | 0.958 | 8, 8 | 1.000 | 0.939 | 1.000 |
| × | × | × | 8, 7 | 0.773 | 0.700 | 0.900 | 8, 7 | 1.000 | 0.750 | 1.000 |

**Table 7.** Mean predictive performance across optimal or near-optimal parameter settings within each model family and each combination of predictors. Performance is measured by the area under the receiver operating characteristic curve and settings that differ in at most 1 parameter by at most 1 grid value are included in the average.

| Perfusate | BAL | Plasma | $n_0, n_3$ | Multiplex | | | Multiplex and Clinical | | | |
| --- | --- | --- | --- | --- | --- | --- | --- | --- | --- | --- |
| | | | | LR | NN | RF | $n_0, n_3$ | LR | NN | RF |
|  |  |  |  |  |  |  | 10, 9 | 0.898 | 0.836 | 0.924 |
| × |  |  | 8, 7 | 0.540 | 0.495 | 0.348 | 8, 7 | 0.963 | 0.767 | 0.818 |
|  | × |  | 8, 8 | 0.812 | 0.670 | 0.906 | 8, 8 | 0.967 | 0.977 | 0.902 |
|  |  | × | 9, 8 | 0.785 | 0.309 | 0.574 | 9, 8 | 0.962 | 0.769 | 0.969 |
| × | × |  | 8, 7 | 0.773 | 0.645 | 0.872 | 8, 7 | 0.955 | 0.704 | 0.735 |
| × |  | × | 8, 7 | 0.435 | 0.200 | 0.425 | 8, 7 | 0.928 | 0.611 | 0.910 |
|  | × | × | 8, 8 | 0.733 | 0.382 | 0.889 | 8, 8 | 0.958 | 0.909 | 0.948 |
| × | × | × | 8, 7 | 0.657 | 0.625 | 0.817 | 8, 7 | 0.971 | 0.711 | 0.904 |

Taken together, these models showed that BAL fluid is the single specimen type with the greatest predictive value. Given this, we built a reduced model using only the BAL data. For each model family (LR and RF), we computed a cumulative measure of the importance of analyte *a* as the sum 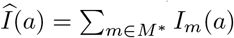of the importance *I*_*m*_(*a*) of *a* in each model in the collection *M* ^∗^ that achieved optimal performance (taken to be zero if *m* did not use *a*). We examined the ranking *a*_1_ ≥ *…* ≥ *a*_71_ of the analytes by cumulative importance and selected the largest number 3≤*k*≤ 8 for which 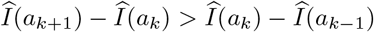 that is, up until a drop-off of importance to the next analyte greater than from the previous. This yielded 6 and 8 analytes from the LR (I-309, Fractalkine, IL-12p40, IL-5, FGF-2, PDGF-AA) and RF (IL-1RA, Fractalkine, M-CSF, IL-6, BCA-1, IP-10, IL-12p40, Eotaxin) model families, respectively, with 2 in common.

We used the most frequent mixture parameter *α*^∗^ = 1 from the models *M* ^∗^ in a regularized regression analysis (hence a LASSO) of these 12 analytes, 4 of which (I-309, IL-5, PDGF-AA, and Eotaxin) were eliminated (Figure 5A). Based on the appearances of the terms and the proportion of deviance explained with increasing penalty, we selected 3 analytes as potentials for a near-patient testing panel capable of disguising between PGD3 or PGD0. Our candidate panel included IL-1RA, Fractalkine, and BCA-1. Figure 5B plots the binary outcome against the logit-transformed probabilities obtained from the model, with the logit curve overlaid to show how these probabilities determine classification. The model accurately predicts every case. A PCA of the 16 cases used to optimize the BAL models on these analytes showed that, together, they clearly separate the PGD subgroups (Figure 5C).

**Figure 5.**
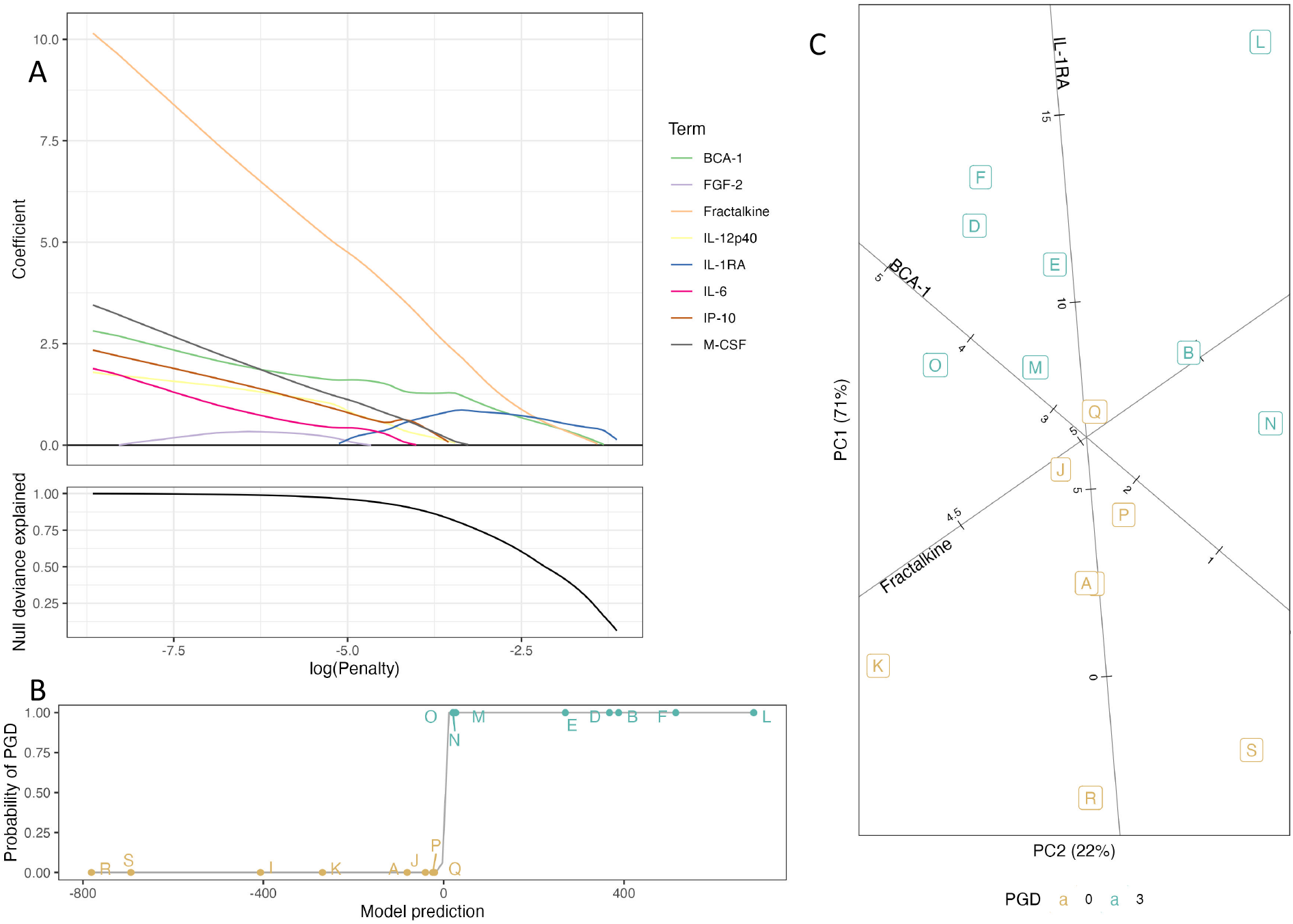
Selecting analytes for a candidate BAL panel. (A) Regularized regression coefficients and proportion of deviance explained in a LASSO on non-imputed multiplex assay data using 8 analytes: BCA-1, FGF-2, Fractalkine, IL-12p40, IL-1RA, IL-6, IP-10, and M-CSF. (B) PGD probability versus linear prediction for 16 cases, derived from a 3-analyte model (BCA-1, Fractalkine, IL-1RA) informed by the LASSO. (C) PCA biplot of 16 cases used in BAL models on the 3 analytes included in the candidate panel.

Because we had exhausted the available cases in the nested CV process used to obtain optimal models, we could not validate this model for performance. Instead, we used classical goodness-of-fit techniques to evaluate the model. While standard diagnostic plots are difficult to interpret for GLMs, those obtained indicated that the reduced model is well-calibrated on most cases, but that a few exceptional cases exert outsized influence. Randomly-generated quantile residuals provide a more appropriate set of diagnostics. We viewed 6 batches, were slightly skewed left but otherwise indistinguishable from Gaussian, which suggests that the model is a reasonable fit to the data. (The issues identified here are to be expected for such a small set of cases.)

### 3.7 Robustness checks

To enhance the rigor of our approach we performed robustness checks. In the main analysis, we utilized values based on protein concentrations, with imputations for exploratory analysis and without imputations for predictive modeling. For robustness checks, we used concentrations without imputations for exploratory analysis and with imputations for predictive modeling, and we further used mean fluorescent intensity values for both. Our checks were qualitatively similar to those obtained in the main analysis. Additionally, optimal performances in the robustness check using FIs were mixed in their comparisons to corresponding performances in the main analysis.

## 4 Discussion

Here, we interrogated the cytokine, chemokine, growth and fibroproliferative factor profiles of multiple biological samples collected from lung transplant recipients for their predictive ability in identifying early PGD development. Our expansive multiplex approach allowed us to examine previously-reported markers of PGD while achieving unbiased, “hypothesis free” discovery of novel biomarkers. Sampling multiple biological sites from each recipient and obtaining results using the same assay and data analysis methodology allowed us to compare the utility of each sample site against each other without bias from batch variability. To further stratify our analysis to patients most at risk, we were cognizant of our inclusion criteria and chose to only examine patients that maintained a PGD grade of 0 or 3 throughout the first 72-hr time course.

To our knowledge, no human studies have investigated cytokine levels in donor lung perfusate collected from vasculature at time of reperfusion. We hypothesized this sample would give us a unique view into static storage and perioperative lung characteristics. Our results show significantly lower levels of IL-17a in perfusate from patients that developed PGD vs those that remained PGD-free. IL-17a is associated with neutrophil chemoattraction/activation and tissue remodeling House et al. (2023). It is therefore unclear if the reduced expression of perfusate IL-17a predisposes PGD+ patients to delayed neutrophil chemoattraction or if another unknown mechanism is at play. Beyond IL-17a, no other analytes were significantly different, and the predictive modeling power of perfusate alone was the lowest of the three biological samples. Recipient plasma samples collected before and after transplant provided us a view into systemic markers indicative of PGD onset. Analysis of plasma samples at individual time points revealed over 15 analytes were significantly different between PGD+ and PGD– patients. Several of these, including IL-6 and MCP-1, align with plasma biomarkers of PGD+ previously reported, while others are original findings (Hoffman et al. 2009; Bharat et al. 2008; Shah et al. 2012; Bonneau et al. 2022; Moreno et al. 2007). Of note, BCA-1 (CXCL13) and IL-22 were significantly elevated in PGD+ plasma compared to PGD– plasma in a multivariate analysis of all time points analyzed. To our knowledge, these cytokines have not been reported as PGD biomarkers thus far, and may indicate that PGD+ patients have increased early signaling for adaptive immune cells and repair/inflammation responses (Mao et al. 2011; Arshad et al. 2020).

Our multivariate predictive modeling comparisons indicated the preeminent sample to analyze for its capacity to predict PGD onset is BAL fluid collected 2hrs post-transplant (T0). Analysis of T0 BAL fluid provides a window into the local lung microenvironment immediately following transplant. Within this sample, we found significantly elevated levels of IL-1Ra, Fractalkine (CX3CL1), BCA-1, Eotaxin, IL-12p40, IL-6, IP-10, and M-CSF in PGD+ samples compared to PGD–. Using a LASSO analysis to analyze the predictive power of analytes in T0 BAL specimen, we identified 3 cytokines—IL-1Ra, BCA-1, and Fractalkine—as strong candidates to accurately predict PGD development while minimizing model deviance. Interestingly, elevated levels of BCA-1 (which we also report in PGD+ plasma) may indicate that increased B lymphocyte activation and signaling occurs with increasing PGD severity, although further work is needed to evaluate this hypothesis (Legler et al. 1998; Jenh et al. 2001). Fractalkine has been previously established as a potential marker of prognosis following ischemic injury in stroke and cardiac transplant (Grosse et al. 2014; Kanzawa et al. 2022; Loh et al. 2023). These studies suggest Fractalkine recruitment of CX3CR1+ leukocytes may contribute to vascular injury (Umehara et al. 2001). Our findings merit the need for studies investigating the role of Fractalkine in post-transplant acute lung injury. Finally, IL-1Ra (Interleukin 1 receptor antagonist) is known to decrease pro-inflammatory signaling and the recruitment of innate immune cells by competitively binding to IL-1R (Arend 2002; Griffith et al. 2023). Additional functional studies on the roles of IL-1Ra, BCA-1, and Fractalkine are needed to fully evaluate their role in PGD severity.

While most biomarker discovery studies use only statistical association tests or univariable prediction models to identify candidates, a growing number use multivariable models, often internally cross-validated (Halloran et al. 2022). For example, Wolf et al. (2011) put a panel of 7 BOS-discriminating proteins obtained from mass spectrometry of BAL fluid into a RF classifier that successfully risk-stratified patients both alone and in combination with FEV1, and Berra et al. (2021) built several ML CLAD prediction models of varying performance from the Western blot–measured concentrations of several peptides in BAL fluid. No multivariable studies have utilized patient-matched biospecimens from different sites—perfusate, BAL fluid, and plasma. While some composite markers have been adapted to candidate point-of-care tests, this has not been done for protein panels (de Silva et al. 2022), and most ML-driven biomarker discovery does not progress to the clinical validation stage (Glaab et al. 2021). Thus, our approach brings high potential for point-of-care testing to the field.

Our analysis found the most significant changes between PGD+ and PGD– patient groups in BAL specimens collected at T0 and suggested that 3 key analytes could be used to accurate predictive the likelihood of PGD onset. Combining donor and recipient clinical data with this BAL data yielded still stronger predictive performance. While our results stood firm in the face of multiple data robustness checks, it is crucial that large, multi-center studies are performed to identify the most accurate analytes to drive predictive modeling, as well as to ensure full representation of diverse patient populations.

Following additional studies, these results have potential for integration into a clinical point-of-care assessment such that recipient BAL fluid may be tested for specific analytes immediately post-transplant as a marker of PGD development and/or severity (i.e. BCA-1, Fractalkine, IL-1Ra). A rapid test of this nature could assist clinicians in managing and stratifying care. Further including factors such as clinical and donor characteristics could strengthen predictive models further, as we have shown in this study. Overall, our results provide basis for a tool to supplement PGD diagnostic criteria and stratify patients by degree of acute lung injury. Additionally, as the need to accurately grade and stratify PGD severity becomes increasingly complex, our proposed predictive modeling approach offers a solution to further classify patients based off relevant biological biomarkers.

### 4.1 Machine learning versus statistical testing for predictor selection

For selecting PGD subgroup discriminants, the ML results partly corroborate the statistical results: While they have the majority of analytes in common, the differences speak to the limitations of both approaches.

Most predictors obtained from variable importance filtering with BAL fluid (BCA-1, Eotaxin, Fractalkine, IL-1RA, and M-CSF) are among those identified by KW tests with a *p <* 0.05 threshold. Of the remainder, two (Eotaxin-3 and TPO) consist almost entirely of imputed values, and all three (including IL-22) are useful only because of one or two outliers (Figure 4C); as such, we manually excluded them as panel candidates. Conversely, the ML approach excluded some discriminants detected by KW tests (IL-12p40, IL-6, IP-10). These might have been excluded by a correlation analysis or stepwise model selection process, but these would have relied further on null hypothesis testing, whereas the ML approach relies only on predictive performance.

## Supporting information

Methodological Supplement

## 5 Availability statement

Matched assay and clinical data are being prepared for deposition to a public repository. All code used in computations and manuscript preparation will be made publicly available upon acceptance for publication.

## 6 Acknowledgments

This work was supported by the National Heart Lung and Blood Institute of the National Institutes of Health under award number 5R01HL140470 (CA).

## Notes

### Competing Interest Statement

The authors have declared no competing interest.

### Summary of Updates

This revision acknowledges funding support omitted in the original version.

