## Supplementary material for "Predicting Primary Graft Dysfunction in Lung Transplantation: Machine Learning–Guided Biomarker Discovery": Methodological Supplement

### Methodological Supplement for: Predicting Primary Graft Dysfunction in Lung Transplantation–Machine Learning–Guided Biomarker Discovery

2024 May 22

**Abstract**

This supplemental text explains the methodology of the study in more detail.

#### Preprocessing

##### Donor organ data

Donor age, gender, race, and smoking and alcohol history were collected. Donor smoking and alcohol history were recoded to yes/no responses. All organs were procured after brain death. Storage temperature was coded as standard or normothermic.

##### Assay data

Plasma samples were taken at 6 time points, coded as "pre" (specimens taken within 12hr before transplantation), "2hr" (specimens taken within 2 hours following transplantation, designated in the text as "T0"), "24hr", "72hr", "1wk", or "2wk" (specimens taken at 14 or 15 days). We used only provided average values in our analysis.

Samples were collected and analyzed from 17 cases (9 PGD–, 8 PGD+). However, multiplex data from perfusate, BAL fluid, and plasma at 2wk were not obtained for an additional PGD– case, from plasma at 2wk for an additional PGD– case, and from perfusate for an additional PGD+ case. While only 13 cases had complete data, analyses using data from only some sites included 15 cases or more. Table M1 reports completeness rates for all proteins.

Table M1: Number and percent of cases for which concentration values were calculatable for each protein.

| Medium | Time | Protein | PGD– | PGD+ |
|---|---|---|---|---|
| perfusate | pre | 6CKine | 6/8 (75.0%) | 7/7 (100.0%) |
| perfusate | pre | EGF | 3/8 (37.5%) | 3/7 (42.9%) |
| perfusate | pre | ENA-78 | 7/8 (87.5%) | 7/7 (100.0%) |
| perfusate | pre | Eotaxin-3 | 3/8 (37.5%) | 2/7 (28.6%) |
| perfusate | pre | Fractalkine | 7/8 (87.5%) | 7/7 (100.0%) |
| perfusate | pre | GM-CSF | 6/8 (75.0%) | 6/7 (85.7%) |
| perfusate | pre | IFN-α2 | 7/8 (87.5%) | 7/7 (100.0%) |
| perfusate | pre | IFNγ | 1/8 (12.5%) | 0/7 (0.0%) |
| perfusate | pre | IL-12p70 | 5/8 (62.5%) | 6/7 (85.7%) |
| perfusate | pre | IL-17A | 6/8 (75.0%) | 4/7 (57.1%) |
| perfusate | pre | IL-17E/IL-25 | 0/8 (0.0%) | 1/7 (14.3%) |
| perfusate | pre | IL-17F | 5/8 (62.5%) | 4/7 (57.1%) |
| perfusate | pre | IL-1β | 7/8 (87.5%) | 7/7 (100.0%) |
| perfusate | pre | IL-2 | 5/8 (62.5%) | 3/7 (42.9%) |

| Medium | Time | Protein | PGD– | PGD+ |
|---|---|---|---|---|
| perfusate | pre | IL-20 | 0/8 (0.0%) | 0/7 (0.0%) |
| perfusate | pre | IL-21 | 0/8 (0.0%) | 0/7 (0.0%) |
| perfusate | pre | IL-22 | 1/8 (12.5%) | 0/7 (0.0%) |
| perfusate | pre | IL-23 | 5/8 (62.5%) | 3/7 (42.9%) |
| perfusate | pre | IL-27 | 7/8 (87.5%) | 5/7 (71.4%) |
| perfusate | pre | IL-28A | 2/8 (25.0%) | 3/7 (42.9%) |
| perfusate | pre | IL-3 | 0/8 (0.0%) | 0/7 (0.0%) |
| perfusate | pre | IL-33 | 7/8 (87.5%) | 7/7 (100.0%) |
| perfusate | pre | IL-4 | 1/8 (12.5%) | 1/7 (14.3%) |
| perfusate | pre | IL-7 | 1/8 (12.5%) | 4/7 (57.1%) |
| perfusate | pre | IL-9 | 7/8 (87.5%) | 7/7 (100.0%) |
| perfusate | pre | LIF | 0/8 (0.0%) | 0/7 (0.0%) |
| perfusate | pre | MCP-2 | 4/8 (50.0%) | 3/7 (42.9%) |
| perfusate | pre | MCP-4 | 2/8 (25.0%) | 0/7 (0.0%) |
| perfusate | pre | MIP-1α | 1/8 (12.5%) | 1/7 (14.3%) |
| perfusate | pre | TNFβ | 2/8 (25.0%) | 2/7 (28.6%) |
| perfusate | pre | TPO | 6/8 (75.0%) | 6/7 (85.7%) |
| perfusate | pre | TSLP | 0/8 (0.0%) | 0/7 (0.0%) |
| perfusate | pre | VEGF-A | 1/8 (12.5%) | 0/7 (0.0%) |
| plasma | pre | 6CKine | 8/9 (88.9%) | 6/8 (75.0%) |
| plasma | pre | EGF | 7/9 (77.8%) | 5/8 (62.5%) |
| plasma | pre | ENA-78 | 8/9 (88.9%) | 8/8 (100.0%) |
| plasma | pre | Eotaxin-3 | 2/9 (22.2%) | 2/8 (25.0%) |
| plasma | pre | FLT-3L | 9/9 (100.0%) | 7/8 (87.5%) |
| plasma | pre | GM-CSF | 8/9 (88.9%) | 7/8 (87.5%) |
| plasma | pre | GROα | 9/9 (100.0%) | 6/8 (75.0%) |
| plasma | pre | IFN-α2 | 6/9 (66.7%) | 6/8 (75.0%) |
| plasma | pre | IFNγ | 7/9 (77.8%) | 8/8 (100.0%) |
| plasma | pre | IL-10 | 4/9 (44.4%) | 6/8 (75.0%) |
| plasma | pre | IL-12p40 | 5/9 (55.6%) | 6/8 (75.0%) |
| plasma | pre | IL-12p70 | 7/9 (77.8%) | 7/8 (87.5%) |
| plasma | pre | IL-13 | 5/9 (55.6%) | 6/8 (75.0%) |
| plasma | pre | IL-15 | 3/9 (33.3%) | 6/8 (75.0%) |
| plasma | pre | IL-16 | 7/9 (77.8%) | 5/8 (62.5%) |
| plasma | pre | IL-17A | 7/9 (77.8%) | 4/8 (50.0%) |
| plasma | pre | IL-17F | 9/9 (100.0%) | 7/8 (87.5%) |
| plasma | pre | IL-2 | 2/9 (22.2%) | 3/8 (37.5%) |
| plasma | pre | IL-20 | 2/9 (22.2%) | 0/8 (0.0%) |
| plasma | pre | IL-21 | 3/9 (33.3%) | 1/8 (12.5%) |
| plasma | pre | IL-22 | 3/9 (33.3%) | 6/8 (75.0%) |
| plasma | pre | IL-23 | 3/9 (33.3%) | 2/8 (25.0%) |
| plasma | pre | IL-28A | 1/9 (11.1%) | 1/8 (12.5%) |
| plasma | pre | IL-3 | 4/9 (44.4%) | 3/8 (37.5%) |
| plasma | pre | IL-33 | 3/9 (33.3%) | 2/8 (25.0%) |
| plasma | pre | IL-7 | 8/9 (88.9%) | 7/8 (87.5%) |
| plasma | pre | IL-9 | 3/9 (33.3%) | 3/8 (37.5%) |
| plasma | pre | LIF | 5/9 (55.6%) | 5/8 (62.5%) |
| plasma | pre | M-CSF | 9/9 (100.0%) | 6/8 (75.0%) |
| plasma | pre | MCP-4 | 9/9 (100.0%) | 6/8 (75.0%) |
| plasma | pre | MIP-1α | 7/9 (77.8%) | 5/8 (62.5%) |
| plasma | pre | SCF | 4/9 (44.4%) | 5/8 (62.5%) |
| plasma | pre | TGFα | 8/9 (88.9%) | 5/8 (62.5%) |

| Medium | Time | Protein | PGD– | PGD+ |
|---|---|---|---|---|
| plasma | pre | TNFβ | 2/9 (22.2%) | 3/8 (37.5%) |
| plasma | pre | TPO | 6/9 (66.7%) | 6/8 (75.0%) |
| plasma | pre | TSLP | 3/9 (33.3%) | 3/8 (37.5%) |
| plasma | 2hr | EGF | 5/9 (55.6%) | 7/8 (87.5%) |
| plasma | 2hr | Eotaxin-2 | 9/9 (100.0%) | 7/8 (87.5%) |
| plasma | 2hr | Eotaxin-3 | 2/9 (22.2%) | 0/8 (0.0%) |
| plasma | 2hr | FLT-3L | 8/9 (88.9%) | 8/8 (100.0%) |
| plasma | 2hr | IFN-α2 | 8/9 (88.9%) | 7/8 (87.5%) |
| plasma | 2hr | IFNγ | 8/9 (88.9%) | 8/8 (100.0%) |
| plasma | 2hr | IL-12p40 | 6/9 (66.7%) | 5/8 (62.5%) |
| plasma | 2hr | IL-13 | 6/9 (66.7%) | 8/8 (100.0%) |
| plasma | 2hr | IL-16 | 6/9 (66.7%) | 7/8 (87.5%) |
| plasma | 2hr | IL-17A | 8/9 (88.9%) | 8/8 (100.0%) |
| plasma | 2hr | IL-17F | 8/9 (88.9%) | 8/8 (100.0%) |
| plasma | 2hr | IL-2 | 6/9 (66.7%) | 8/8 (100.0%) |
| plasma | 2hr | IL-20 | 0/9 (0.0%) | 0/8 (0.0%) |
| plasma | 2hr | IL-21 | 1/9 (11.1%) | 1/8 (12.5%) |
| plasma | 2hr | IL-22 | 7/9 (77.8%) | 8/8 (100.0%) |
| plasma | 2hr | IL-23 | 1/9 (11.1%) | 1/8 (12.5%) |
| plasma | 2hr | IL-28A | 1/9 (11.1%) | 0/8 (0.0%) |
| plasma | 2hr | IL-3 | 5/9 (55.6%) | 3/8 (37.5%) |
| plasma | 2hr | IL-33 | 1/9 (11.1%) | 1/8 (12.5%) |
| plasma | 2hr | IL-7 | 7/9 (77.8%) | 8/8 (100.0%) |
| plasma | 2hr | IL-9 | 2/9 (22.2%) | 2/8 (25.0%) |
| plasma | 2hr | LIF | 6/9 (66.7%) | 5/8 (62.5%) |
| plasma | 2hr | MCP-4 | 6/9 (66.7%) | 2/8 (25.0%) |
| plasma | 2hr | SCF | 5/9 (55.6%) | 3/8 (37.5%) |
| plasma | 2hr | TGFα | 8/9 (88.9%) | 8/8 (100.0%) |
| plasma | 2hr | TNFβ | 3/9 (33.3%) | 2/8 (25.0%) |
| plasma | 2hr | TPO | 4/9 (44.4%) | 2/8 (25.0%) |
| plasma | 2hr | TSLP | 1/9 (11.1%) | 1/8 (12.5%) |
| plasma | 2hr | VEGF-A | 9/9 (100.0%) | 7/8 (87.5%) |
| plasma | 24hr | 6CKine | 8/9 (88.9%) | 8/8 (100.0%) |
| plasma | 24hr | EGF | 6/9 (66.7%) | 6/8 (75.0%) |
| plasma | 24hr | ENA-78 | 7/9 (77.8%) | 8/8 (100.0%) |
| plasma | 24hr | Eotaxin-3 | 0/9 (0.0%) | 0/8 (0.0%) |
| plasma | 24hr | FLT-3L | 8/9 (88.9%) | 7/8 (87.5%) |
| plasma | 24hr | GM-CSF | 8/9 (88.9%) | 8/8 (100.0%) |
| plasma | 24hr | GROα | 6/9 (66.7%) | 8/8 (100.0%) |
| plasma | 24hr | IFN-α2 | 6/9 (66.7%) | 5/8 (62.5%) |
| plasma | 24hr | IL-12p40 | 4/9 (44.4%) | 2/8 (25.0%) |
| plasma | 24hr | IL-12p70 | 8/9 (88.9%) | 7/8 (87.5%) |
| plasma | 24hr | IL-13 | 6/9 (66.7%) | 6/8 (75.0%) |
| plasma | 24hr | IL-16 | 8/9 (88.9%) | 7/8 (87.5%) |
| plasma | 24hr | IL-17A | 6/9 (66.7%) | 6/8 (75.0%) |
| plasma | 24hr | IL-17E/IL-25 | 9/9 (100.0%) | 7/8 (87.5%) |
| plasma | 24hr | IL-17F | 6/9 (66.7%) | 3/8 (37.5%) |
| plasma | 24hr | IL-20 | 0/9 (0.0%) | 0/8 (0.0%) |
| plasma | 24hr | IL-21 | 1/9 (11.1%) | 0/8 (0.0%) |
| plasma | 24hr | IL-22 | 6/9 (66.7%) | 8/8 (100.0%) |
| plasma | 24hr | IL-23 | 1/9 (11.1%) | 1/8 (12.5%) |
| plasma | 24hr | IL-28A | 1/9 (11.1%) | 0/8 (0.0%) |

| Medium | Time | Protein | PGD– | PGD+ |
|---|---|---|---|---|
| plasma | 24hr | IL-3 | 4/9 (44.4%) | 3/8 (37.5%) |
| plasma | 24hr | IL-33 | 1/9 (11.1%) | 1/8 (12.5%) |
| plasma | 24hr | IL-7 | 8/9 (88.9%) | 8/8 (100.0%) |
| plasma | 24hr | IL-9 | 2/9 (22.2%) | 2/8 (25.0%) |
| plasma | 24hr | LIF | 3/9 (33.3%) | 4/8 (50.0%) |
| plasma | 24hr | MCP-4 | 4/9 (44.4%) | 2/8 (25.0%) |
| plasma | 24hr | MIP-1α | 7/9 (77.8%) | 7/8 (87.5%) |
| plasma | 24hr | SCF | 3/9 (33.3%) | 2/8 (25.0%) |
| plasma | 24hr | TGFα | 7/9 (77.8%) | 7/8 (87.5%) |
| plasma | 24hr | TNFβ | 2/9 (22.2%) | 0/8 (0.0%) |
| plasma | 24hr | TPO | 5/9 (55.6%) | 3/8 (37.5%) |
| plasma | 24hr | TRAIL | 9/9 (100.0%) | 7/8 (87.5%) |
| plasma | 24hr | TSLP | 1/9 (11.1%) | 1/8 (12.5%) |
| plasma | 72hr | 6CKine | 9/9 (100.0%) | 7/8 (87.5%) |
| plasma | 72hr | EGF | 7/9 (77.8%) | 5/8 (62.5%) |
| plasma | 72hr | ENA-78 | 9/9 (100.0%) | 7/8 (87.5%) |
| plasma | 72hr | Eotaxin-3 | 2/9 (22.2%) | 1/8 (12.5%) |
| plasma | 72hr | FLT-3L | 9/9 (100.0%) | 6/8 (75.0%) |
| plasma | 72hr | GROα | 6/9 (66.7%) | 6/8 (75.0%) |
| plasma | 72hr | IFN-α2 | 7/9 (77.8%) | 5/8 (62.5%) |
| plasma | 72hr | IFNγ | 8/9 (88.9%) | 8/8 (100.0%) |
| plasma | 72hr | IL-12p40 | 5/9 (55.6%) | 2/8 (25.0%) |
| plasma | 72hr | IL-12p70 | 9/9 (100.0%) | 6/8 (75.0%) |
| plasma | 72hr | IL-13 | 6/9 (66.7%) | 6/8 (75.0%) |
| plasma | 72hr | IL-16 | 4/9 (44.4%) | 6/8 (75.0%) |
| plasma | 72hr | IL-17A | 7/9 (77.8%) | 6/8 (75.0%) |
| plasma | 72hr | IL-17E/IL-25 | 9/9 (100.0%) | 7/8 (87.5%) |
| plasma | 72hr | IL-17F | 8/9 (88.9%) | 6/8 (75.0%) |
| plasma | 72hr | IL-2 | 7/9 (77.8%) | 7/8 (87.5%) |
| plasma | 72hr | IL-20 | 0/9 (0.0%) | 0/8 (0.0%) |
| plasma | 72hr | IL-21 | 1/9 (11.1%) | 1/8 (12.5%) |
| plasma | 72hr | IL-22 | 6/9 (66.7%) | 6/8 (75.0%) |
| plasma | 72hr | IL-23 | 2/9 (22.2%) | 3/8 (37.5%) |
| plasma | 72hr | IL-28A | 1/9 (11.1%) | 0/8 (0.0%) |
| plasma | 72hr | IL-3 | 7/9 (77.8%) | 2/8 (25.0%) |
| plasma | 72hr | IL-33 | 1/9 (11.1%) | 2/8 (25.0%) |
| plasma | 72hr | IL-5 | 8/9 (88.9%) | 6/8 (75.0%) |
| plasma | 72hr | IL-7 | 8/9 (88.9%) | 7/8 (87.5%) |
| plasma | 72hr | IL-9 | 6/9 (66.7%) | 2/8 (25.0%) |
| plasma | 72hr | LIF | 7/9 (77.8%) | 7/8 (87.5%) |
| plasma | 72hr | MCP-2 | 9/9 (100.0%) | 7/8 (87.5%) |
| plasma | 72hr | MCP-4 | 5/9 (55.6%) | 2/8 (25.0%) |
| plasma | 72hr | MIP-1α | 8/9 (88.9%) | 7/8 (87.5%) |
| plasma | 72hr | SCF | 3/9 (33.3%) | 2/8 (25.0%) |
| plasma | 72hr | TNFβ | 3/9 (33.3%) | 0/8 (0.0%) |
| plasma | 72hr | TPO | 5/9 (55.6%) | 6/8 (75.0%) |
| plasma | 72hr | TSLP | 1/9 (11.1%) | 3/8 (37.5%) |
| plasma | 1wk | EGF | 6/9 (66.7%) | 5/8 (62.5%) |
| plasma | 1wk | ENA-78 | 8/9 (88.9%) | 7/8 (87.5%) |
| plasma | 1wk | Eotaxin-3 | 1/9 (11.1%) | 0/8 (0.0%) |
| plasma | 1wk | GROα | 8/9 (88.9%) | 7/8 (87.5%) |
| plasma | 1wk | IFN-α2 | 7/9 (77.8%) | 6/8 (75.0%) |

| Medium | Time | Protein | PGD– | PGD+ |
| --- | --- | --- | --- | --- |
| plasma | 1wk | IL-12p40 | 4/9 (44.4%) | 2/8 (25.0%) |
| plasma | 1wk | IL-12p70 | 9/9 (100.0%) | 7/8 (87.5%) |
| plasma | 1wk | IL-13 | 7/9 (77.8%) | 7/8 (87.5%) |
| plasma | 1wk | IL-16 | 6/9 (66.7%) | 6/8 (75.0%) |
| plasma | 1wk | IL-17A | 6/9 (66.7%) | 7/8 (87.5%) |
| plasma | 1wk | IL-17E/IL-25 | 8/9 (88.9%) | 7/8 (87.5%) |
| plasma | 1wk | IL-17F | 7/9 (77.8%) | 6/8 (75.0%) |
| plasma | 1wk | IL-2 | 7/9 (77.8%) | 7/8 (87.5%) |
| plasma | 1wk | IL-20 | 1/9 (11.1%) | 0/8 (0.0%) |
| plasma | 1wk | IL-21 | 0/9 (0.0%) | 0/8 (0.0%) |
| plasma | 1wk | IL-22 | 5/9 (55.6%) | 7/8 (87.5%) |
| plasma | 1wk | IL-23 | 3/9 (33.3%) | 1/8 (12.5%) |
| plasma | 1wk | IL-28A | 0/9 (0.0%) | 0/8 (0.0%) |
| plasma | 1wk | IL-3 | 6/9 (66.7%) | 3/8 (37.5%) |
| plasma | 1wk | IL-33 | 2/9 (22.2%) | 1/8 (12.5%) |
| plasma | 1wk | IL-5 | 9/9 (100.0%) | 7/8 (87.5%) |
| plasma | 1wk | IL-7 | 8/9 (88.9%) | 8/8 (100.0%) |
| plasma | 1wk | IL-9 | 4/9 (44.4%) | 2/8 (25.0%) |
| plasma | 1wk | LIF | 7/9 (77.8%) | 5/8 (62.5%) |
| plasma | 1wk | MCP-2 | 9/9 (100.0%) | 7/8 (87.5%) |
| plasma | 1wk | MCP-4 | 8/9 (88.9%) | 4/8 (50.0%) |
| plasma | 1wk | MIP-1α | 8/9 (88.9%) | 6/8 (75.0%) |
| plasma | 1wk | SCF | 5/9 (55.6%) | 7/8 (87.5%) |
| plasma | 1wk | TGFα | 9/9 (100.0%) | 7/8 (87.5%) |
| plasma | 1wk | TNFβ | 2/9 (22.2%) | 2/8 (25.0%) |
| plasma | 1wk | TPO | 4/9 (44.4%) | 5/8 (62.5%) |
| plasma | 1wk | TSLP | 2/9 (22.2%) | 3/8 (37.5%) |
| plasma | 2wk | 6CKine | 7/7 (100.0%) | 6/7 (85.7%) |
| plasma | 2wk | EGF | 2/7 (28.6%) | 4/7 (57.1%) |
| plasma | 2wk | ENA-78 | 4/7 (57.1%) | 6/7 (85.7%) |
| plasma | 2wk | Eotaxin-3 | 1/7 (14.3%) | 0/7 (0.0%) |
| plasma | 2wk | FLT-3L | 7/7 (100.0%) | 6/7 (85.7%) |
| plasma | 2wk | GROα | 2/7 (28.6%) | 5/7 (71.4%) |
| plasma | 2wk | IFN-α2 | 6/7 (85.7%) | 5/7 (71.4%) |
| plasma | 2wk | IFNγ | 5/7 (71.4%) | 6/7 (85.7%) |
| plasma | 2wk | IL-12p40 | 3/7 (42.9%) | 1/7 (14.3%) |
| plasma | 2wk | IL-12p70 | 7/7 (100.0%) | 6/7 (85.7%) |
| plasma | 2wk | IL-13 | 5/7 (71.4%) | 6/7 (85.7%) |
| plasma | 2wk | IL-16 | 4/7 (57.1%) | 3/7 (42.9%) |
| plasma | 2wk | IL-17A | 6/7 (85.7%) | 6/7 (85.7%) |
| plasma | 2wk | IL-17E/IL-25 | 7/7 (100.0%) | 4/7 (57.1%) |
| plasma | 2wk | IL-17F | 6/7 (85.7%) | 6/7 (85.7%) |
| plasma | 2wk | IL-2 | 5/7 (71.4%) | 2/7 (28.6%) |
| plasma | 2wk | IL-20 | 0/7 (0.0%) | 0/7 (0.0%) |
| plasma | 2wk | IL-21 | 0/7 (0.0%) | 0/7 (0.0%) |
| plasma | 2wk | IL-22 | 4/7 (57.1%) | 6/7 (85.7%) |
| plasma | 2wk | IL-23 | 1/7 (14.3%) | 1/7 (14.3%) |
| plasma | 2wk | IL-28A | 0/7 (0.0%) | 0/7 (0.0%) |
| plasma | 2wk | IL-3 | 6/7 (85.7%) | 4/7 (57.1%) |
| plasma | 2wk | IL-33 | 0/7 (0.0%) | 0/7 (0.0%) |
| plasma | 2wk | IL-9 | 4/7 (57.1%) | 2/7 (28.6%) |
| plasma | 2wk | LIF | 3/7 (42.9%) | 4/7 (57.1%) |

| Medium | Time | Protein | PGD– | PGD+ |
|---|---|---|---|---|
| plasma | 2wk | MCP-2 | 7/7 (100.0%) | 6/7 (85.7%) |
| plasma | 2wk | MIP-1α | 6/7 (85.7%) | 5/7 (71.4%) |
| plasma | 2wk | SCF | 4/7 (57.1%) | 6/7 (85.7%) |
| plasma | 2wk | TGFα | 6/7 (85.7%) | 6/7 (85.7%) |
| plasma | 2wk | TNFβ | 1/7 (14.3%) | 0/7 (0.0%) |
| plasma | 2wk | TPO | 2/7 (28.6%) | 2/7 (28.6%) |
| plasma | 2wk | TSLP | 0/7 (0.0%) | 2/7 (28.6%) |
| plasma | 2wk | VEGF-A | 6/7 (85.7%) | 7/7 (100.0%) |
| BAL | 2hr | 6CKine | 1/8 (12.5%) | 2/8 (25.0%) |
| BAL | 2hr | CTACK | 6/8 (75.0%) | 8/8 (100.0%) |
| BAL | 2hr | EGF | 8/8 (100.0%) | 6/8 (75.0%) |
| BAL | 2hr | Eotaxin-3 | 1/8 (12.5%) | 1/8 (12.5%) |
| BAL | 2hr | GM-CSF | 7/8 (87.5%) | 6/8 (75.0%) |
| BAL | 2hr | IFNγ | 8/8 (100.0%) | 7/8 (87.5%) |
| BAL | 2hr | IL-17E/IL-25 | 8/8 (100.0%) | 7/8 (87.5%) |
| BAL | 2hr | IL-17F | 7/8 (87.5%) | 7/8 (87.5%) |
| BAL | 2hr | IL-20 | 0/8 (0.0%) | 1/8 (12.5%) |
| BAL | 2hr | IL-21 | 0/8 (0.0%) | 1/8 (12.5%) |
| BAL | 2hr | IL-22 | 4/8 (50.0%) | 6/8 (75.0%) |
| BAL | 2hr | IL-23 | 3/8 (37.5%) | 4/8 (50.0%) |
| BAL | 2hr | IL-27 | 7/8 (87.5%) | 5/8 (62.5%) |
| BAL | 2hr | IL-28A | 0/8 (0.0%) | 1/8 (12.5%) |
| BAL | 2hr | IL-3 | 1/8 (12.5%) | 1/8 (12.5%) |
| BAL | 2hr | IL-33 | 2/8 (25.0%) | 3/8 (37.5%) |
| BAL | 2hr | IL-4 | 7/8 (87.5%) | 4/8 (50.0%) |
| BAL | 2hr | IL-7 | 7/8 (87.5%) | 6/8 (75.0%) |
| BAL | 2hr | LIF | 0/8 (0.0%) | 3/8 (37.5%) |
| BAL | 2hr | MCP-2 | 7/8 (87.5%) | 8/8 (100.0%) |
| BAL | 2hr | MCP-4 | 2/8 (25.0%) | 1/8 (12.5%) |
| BAL | 2hr | MIP-1α | 7/8 (87.5%) | 8/8 (100.0%) |
| BAL | 2hr | PDGF-AB/BB | 3/8 (37.5%) | 5/8 (62.5%) |
| BAL | 2hr | TNFβ | 6/8 (75.0%) | 8/8 (100.0%) |
| BAL | 2hr | TPO | 0/8 (0.0%) | 3/8 (37.5%) |
| BAL | 2hr | TSLP | 0/8 (0.0%) | 0/8 (0.0%) |
| BAL | 2hr | VEGF-A | 8/8 (100.0%) | 5/8 (62.5%) |

Distributions of pooled protein concentrations at each time point were right-skewed; those following the offset log-transformation $x \mapsto \log(x+1)$ (henceforth "log-concentrations") were roughly normal, and these values were used throughout. Research IDs used internally were uniquely coded to capital letters for reporting of results.

We included all calculated observed concentrations as numeric values, including those interpolated and those extrapolated using standard limit curves. Remaining entries were out-of-range ("OOR", "OOR <", or "OOR >") and not calculated from fluorescence intensities (FIs). When the FI was below the smallest dilution on the limit curve, we imputed a concentration value obtained as the smallest value calculated for that protein in that medium, across all cases—and, in the case of plasma, across all time points. Analogously, we imputed the largest calculated value for OOR entries when the FI was above the largest dilution on the limit curve.

Multiplex assay data from perfusate and BAL fluid were pre-processed in the same way as those from plasma. Note that one set of limit curves was provided for each protein in perfusate and in BAL concentration measurements.

Proteins were not excluded *a priori* due to high rates of missingness. Instead, proteins were excluded as needed by each analysis, as explained below.

##### Recipient characteristics

Recipient age, sex, and BMI and LAS score at listing were collected. We also determined whether each recipient was on ECMO pre-LTx and post-operatively and whether they received antithymocyte globulin (ATG) treatment. Spirometry results including PaO2, FiO2, and P/F ratio and the presence of bilateral infiltrates on chest radiograph were recorded at 0hr, 24hr, 48hr, and 72hr.

Each recipient was graded for PGD at 0hr, 24hr, 48hr, and 72hr independently by two pulmonologists according to criteria established by the International Heart and Lung Society (Snell et al. 2017). Only patients with a PGD grade of 0 or 3 for the entire 72hr post-transplant period were included in this study and are referred to as PGD− and PGD+, respectively.

#### Protein concentrations and pairwise associations

We visualized co-concentration patterns using heatmaps of proteins by subjects (perfusate and BAL fluid) or subject–time points (plasma) with row and column dendrogram annotations based on complete-linkage clustering by euclidean distance (Kolde 2019). For each medium, and for plasma at the three time points nearest time of transplant, we rendered heatmaps both for all subjects and separately for PGD− and PGD+ subjects.

We generated triangular correlation heatmaps of proteins with at least one correlate for each set of biological media (Wei et al. 2021). Proteins were ordered according to complete-linkage clustering dendrograms.

To characterize changes in concentrations over time (across all post-LTx time points), we used Kruskal–Wallis tests to identify proteins that had non-constant concentrations over time. Based on these values, we opted for the more restrictive threshold $p < 0.01$ because the proteins that met this threshold were adequate to characterize trajectories. We then performed a principal components analysis (PCA) of all subject–time points (including pre-LTx) in the space of the log-concentrations of these proteins. We generated a row-principal biplot of subject trajectories along the first two PCs, and we overlaid segments connecting consecutive centroids in order to visualize the average trajectory.

#### Discrimination of PGD

We visually compared the log-concentrations in perfusate and in BAL as jitter plots and in plasma as line plots between the PGD− and PGD+ subgroups, in each case stratified (main text) or faceted (Figures M1 and M2) by protein. Facets were ordered by the p-values obtained from Kruskal–Wallis rank sum (perfusate and BAL) or multivariate Cramér (Baringhaus and Franz 2004) (plasma, using all time points) tests of their ability to discriminate between the PGD− and PGD+ subgroups. Kruskal–Wallis and Cramér tests required at least one observation from each PGD group. We only plotted proteins with at least three observations in each PGD group.

We also generated another biplot of subject trajectories in plasma, with markers color-coded and trajectories stratified by PGD group. Additionally, we conducted PCA and generated biplots, again color-coding markers by PGD group, on log-concentrations in perfusate and in BAL fluid.

#### Prediction of PGD

We used logistic regression (LR), random forest (RF), and nearest neighbors (NN) models in a machine learning (ML) workflow to determine whether several sets of predictors could discriminate between PGD− and PGD+ subjects.

We assessed the predictive value of every nonempty subset of 4 data sources: protein levels in the three media (perfusate, BAL fluid, and first three time points of plasma) and the clinical data (which included donor, recipient, and perioperative characteristics). These data sets included the following predictors:

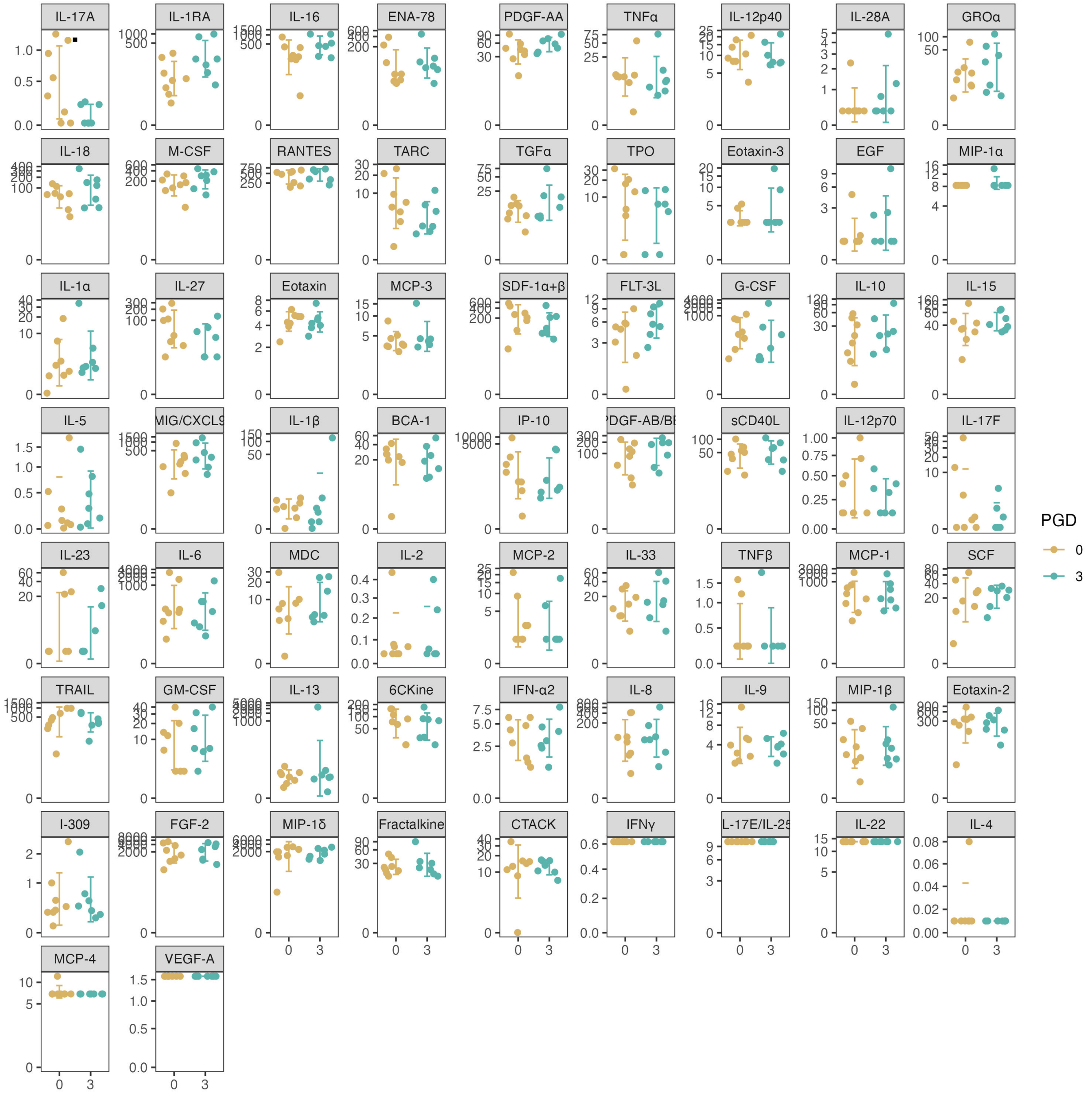


Figure M1: Jitter plots with 1–standard deviation intervals, grouped by PGD grade. The panels are in order of increasing Kruskal–Wallis test p-value and marked as in Figure 2 (main text).

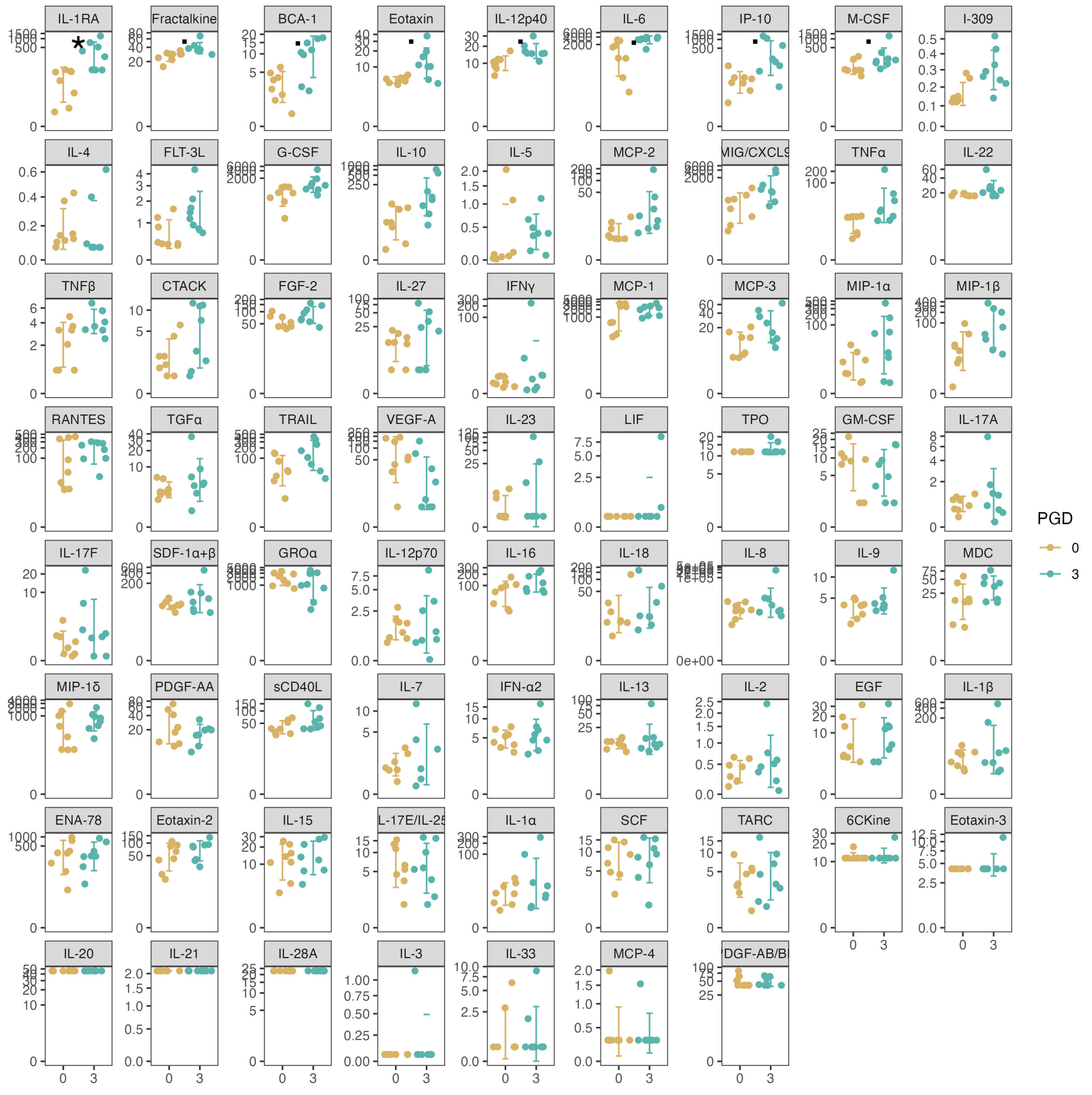


Figure M2: Jitter plots with 1–standard deviation intervals, grouped by PGD grade. The panels are in order of increasing Kruskal–Wallis test p-value and marked as in Figure 2 (main text).

1. Clinical: donor age, donor gender, donor race, storage thermicity, donor smoking history, donor alcohol history, recipient ECMO status pre-Tx and post-op, ATG treatment, recipient LAS score, recipient pre-LTx primary pulmonary diagnosis, recipient age, recipient sex, recipient BMI at listing, and recipient PaO2, FiO2, and P/F ratio for 19 cases.
2. Storage perfusate: multiplex concentrations of 71 proteins at each of 3 time points (pre, 2hr, 24hr) for 16 cases.
3. Post-LTx BAL fluid: multiplex concentrations of 70 proteins (missing IL-7) for 17 cases.
4. Recipient blood plasma: multiplex concentrations of 70 proteins (missing IL-7) for 17 cases.

Combined data were prepared for modeling through a common recipe: All protein concentrations were log-transformed and undefined or infinite values declared missing, and real-valued (including clinical) variables with any missing values were discarded. Categorical (including logical) variables had missing values replaced with the a dummy value for "unknown", while logical variables with complete values were recoded to the integer values 0 and 1. Categorical and logical variables that were constant over our population were discarded. Categorical predictors were expanded into multiple dummy predictors, taking one value of each as the reference value. Finally, any remaining predictors with missing values were discarded.

From the donor, recipient, and perioperative clinical data, organ procurement was discarded due to taking a constant value, and LAS score, recipient BMI, and recipient PaO2, FiO2, and P/F ratio were discarded due to incompleteness. In perfusate, 9 proteins were excluded by the recipe: IFNγ, IL-3, IL-7, IL-22, VEGF-A, IL-20, IL-21, LIF, TSLP. In BAL fluid, 5 proteins were excluded by the recipe: IL-7, IL-20, IL-21, IL-28A, TSLP. Of 213 measurements included from blood plasma over 3 time points, 3 were excluded by the recipe: IL-20 at 2hr, Eotaxin-3 at 24hr, and IL-20 at 24hr.

Models were specified over the following hyperparameter grids:

- LR: penalty $\lambda$ and regularization mixture $\alpha$:

$$\{\lambda = 1 \times 10^{-2\ell} : 0 \leq \ell \leq 5\} \times \{\alpha = a/5 : 0 \leq a \leq 5\}$$

- NN: number of neighbors $k$:

$$\{1 \leq k \leq 9\}$$

- RF: number of trees $\nu$ and maximum feature split $\mu$, where $p$ is the number of predictors:

$$(\{\nu = 1\} \cup \{\nu = 250n : 1 \leq n \leq 8\}) \times \{\mu = \sqrt[3]{p}, \sqrt{p}, p/2, p\}$$

For each model specification, we calculated the area under the receiver operating characteristic curve (AUROC) for leave-one-out cross-validation (LOOCV), stratified by outcome so that each test set comprised one PGD– and one PGD+ case. We then repeated this procedure 12 times to ensure diversity in the test sets. We ranked the model specifications by mean AUROC across 12 LOOCVs.

We summarized the means and standard errors of the AUROCs as line–range plots (Figure M3). We then tabulated the highest AUROC for each combination of data sources and model family. To assess the sensitivity of these optimal performances and present a more realistic expectation of out-of-sample performance, we also tabulated the mean performance of all models whose specifications were within one step on the hyperparameter grid from an optimal specification.

We also examined model-based predictor importance measures for the optimized LR and RF models: absolute coefficient estimates for LR and mean decrease in accuracy for RF. Predictors whose maximum absolute coefficient estimates were smaller than 1% of the largest median absolute coefficient estimate were excluded from the LR evaluation, and those whose mean decrease in accuracy was negative were excluded from the RF evaluation. Predictor importances (using signed coefficient estimates) were ranked by median importance and plotted on cube root–transformed axes for easier visual comparison.

To test the feasibility of a point-of-care test, we extracted the most important predictors from one data set for inclusion in a reduced model.

For each model family (LR and RF), we computed a cumulative measure of the importance of analyte $a$ as the sum $\widehat{I}(a) = \sum_{m \in M^*} I_m(a)$ of the importance $I_m(a)$ of $a$ in each model in the collection $M^*$

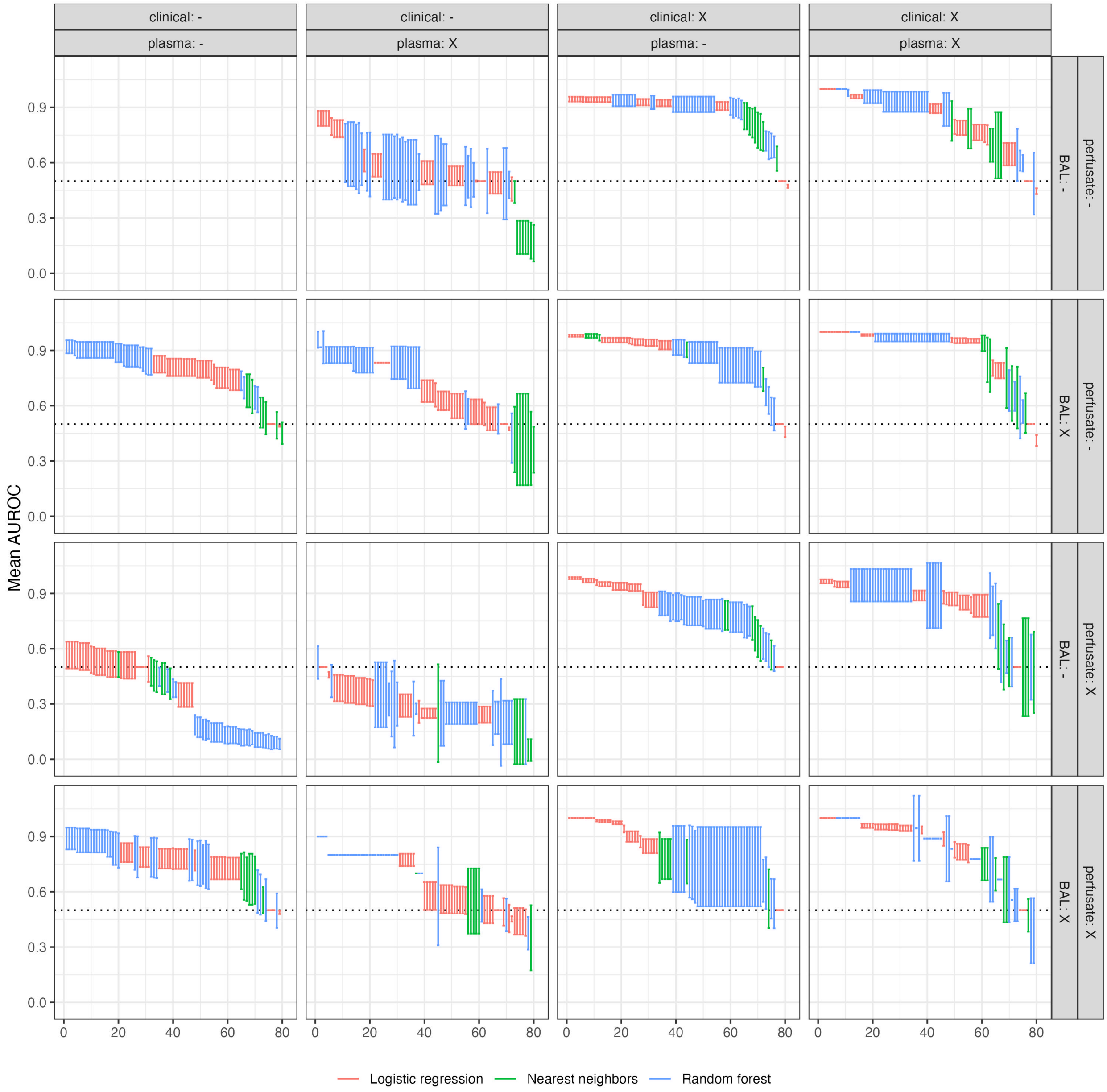


Figure M3: Ranked plots of performances of each model specification across 12 LOOCVs.

that achieved optimal performance (taken to be zero if $m$ did not use $a$). We examined the ranking $a_1 \geq \cdots \geq a_{71}$ of the analytes by cumulative importance and selected the largest number $3 \leq k \leq 8$ for which $\widehat{I}(a_{k+1}) - \widehat{I}(a_k) > \widehat{I}(a_k) - \widehat{I}(a_{k-1})$—that is, up until a drop-off of importance to the next analyte greater than from the previous (Figure M4). This yielded 6 and 8 analytes from the LR (I-309, Fractalkine, IL-12p40, IL-5, FGF-2, PDGF-AA) and RF (IL-1RA, Fractalkine, M-CSF, IL-6, BCA-1, IP-10, IL-12p40, Eotaxin) model families, respectively, with 2 in common.

We pooled the top predictors from the LR and RF models and used LASSO regularization on the full data set to better understand their roles and value in predicting PGD group. Based on the LASSO, we selected a subet as a candidate biomarker panel. We visualized the relationship of the selected predictors to PGD grade using PCA and evaluated the performance and calibration of the logistic regression model using that subset.

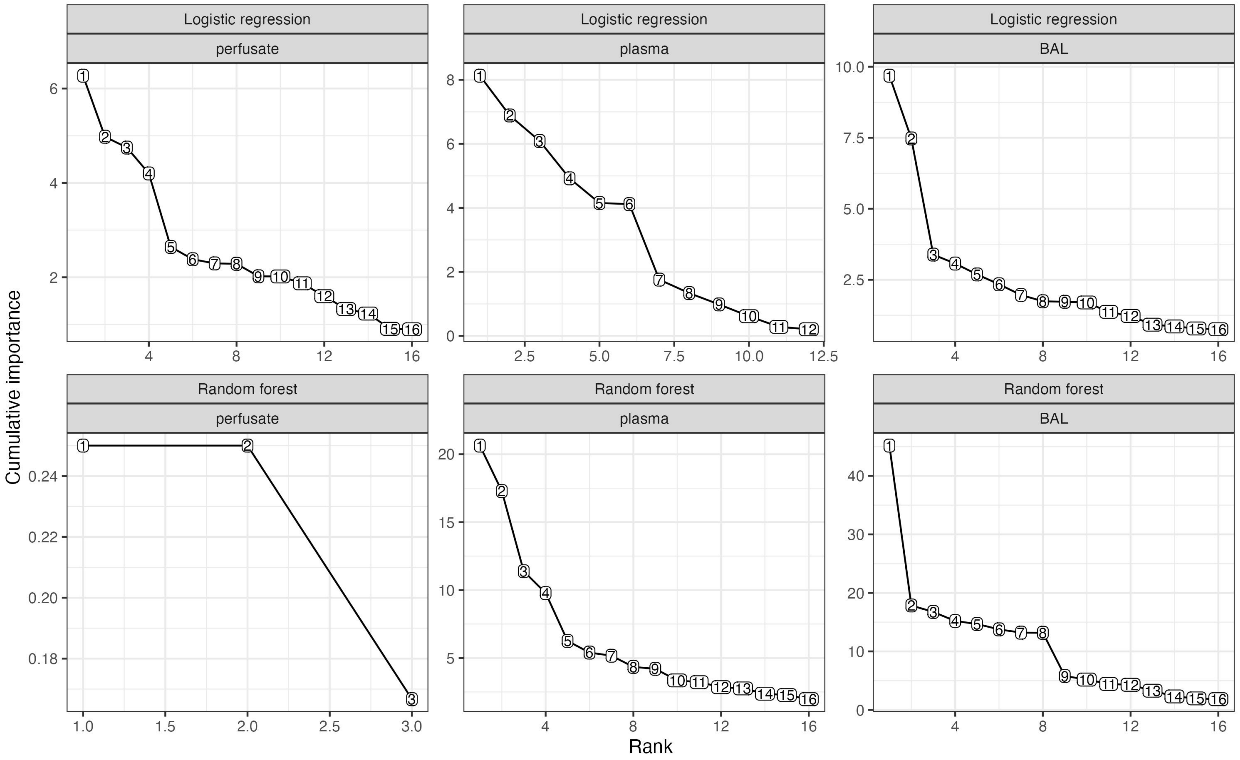


Figure M4: Top 16 cumulative importance measurements $I_C(a)$ for analytes $a$ in each model family, based on optimized models fitted to full data. Cutoffs were imposed at $I_C \leq 5$ (LR) and at $I_C \leq 7$ (RF).

While standard diagnostic plots are difficult to interpret for GLMs, those obtained indicated that the reduced model is well-calibrated on most cases, but that a few exceptional cases exert outsized influence (Figure M5). Randomly-generated quantile residuals (Dunn and Smyth 1996) provide a more appropriate set of diagnostics. We viewed 6 batches, were slightly skewed left but otherwise indistinguishable from Gaussian, which suggests that the model is a reasonable fit to the data (Figure M6). (The issues identified here are to be expected for such a small set of cases.)

#### Robustness checks

In discussion among three authors (JCB, DN, CA), we made three choices regarding the main analysis of multiplex data: First, we chose to include all calculated observed concentrations provided by Eve Technologies, including those extrapolated from outside the limit curves. Second, for exploratory analysis, we chose to impute values where these were missing due to fluorescence intensities being too far outside the limit curve. For values below (above) each limit curve, we imputed the minimum (maximum) calculated observed concentration obtained in the same medium across all cases (and, in the case of plasma, time points). For

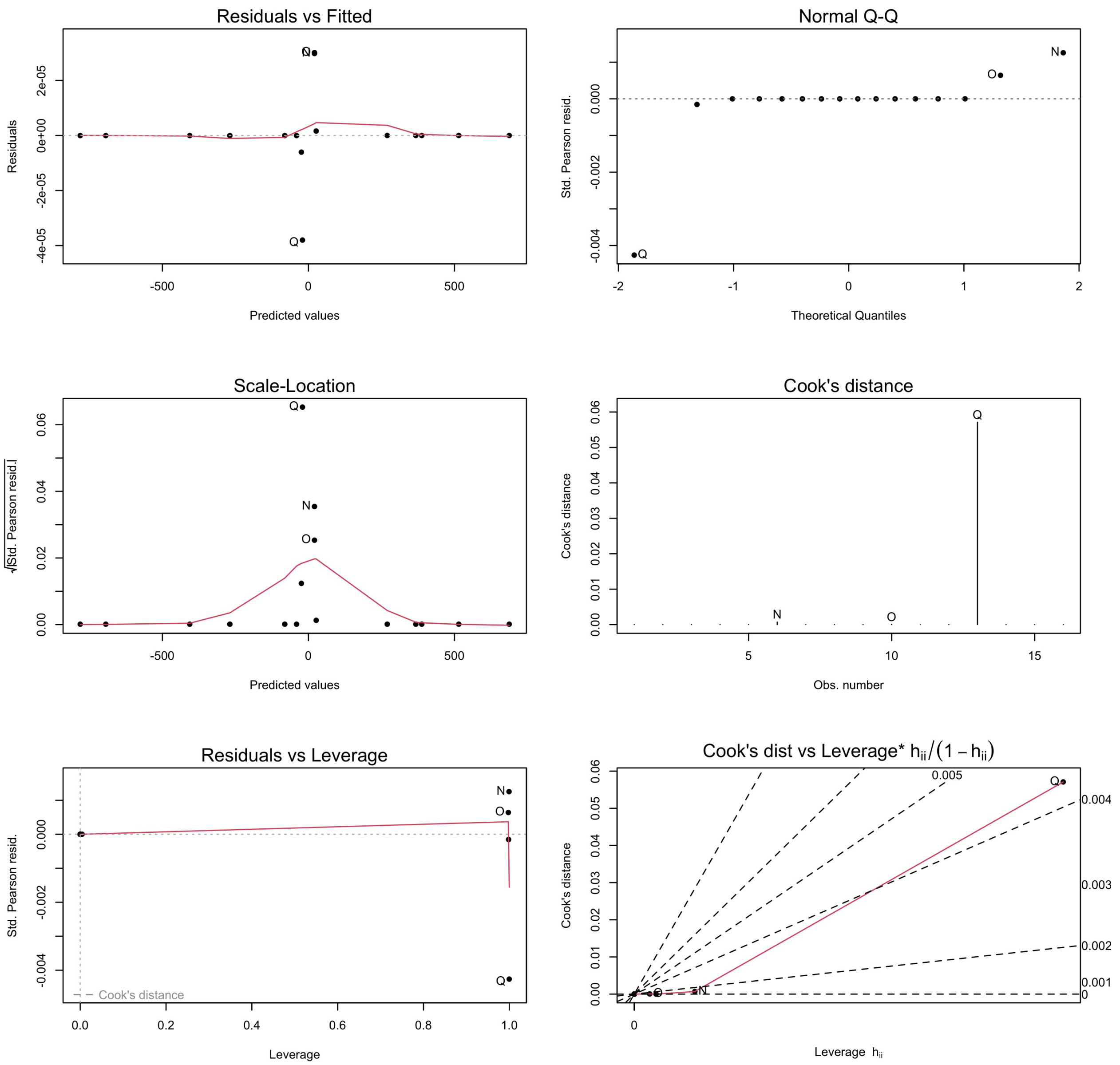


Figure M5: Diagnostic plots for linear regression models provided by ‘plot.lm()‘ in the R stats package; left-to-right, top-to-bottom: (i) residuals versus fitted values, (ii) quantile–quantile normality plot, (iii) root-absolute residuals versus fitted values, (iv) Cook’s distance (versus row index), (v) residuals versus leverage, and (vi) Cook’s distance versus odds-transformed leverage.

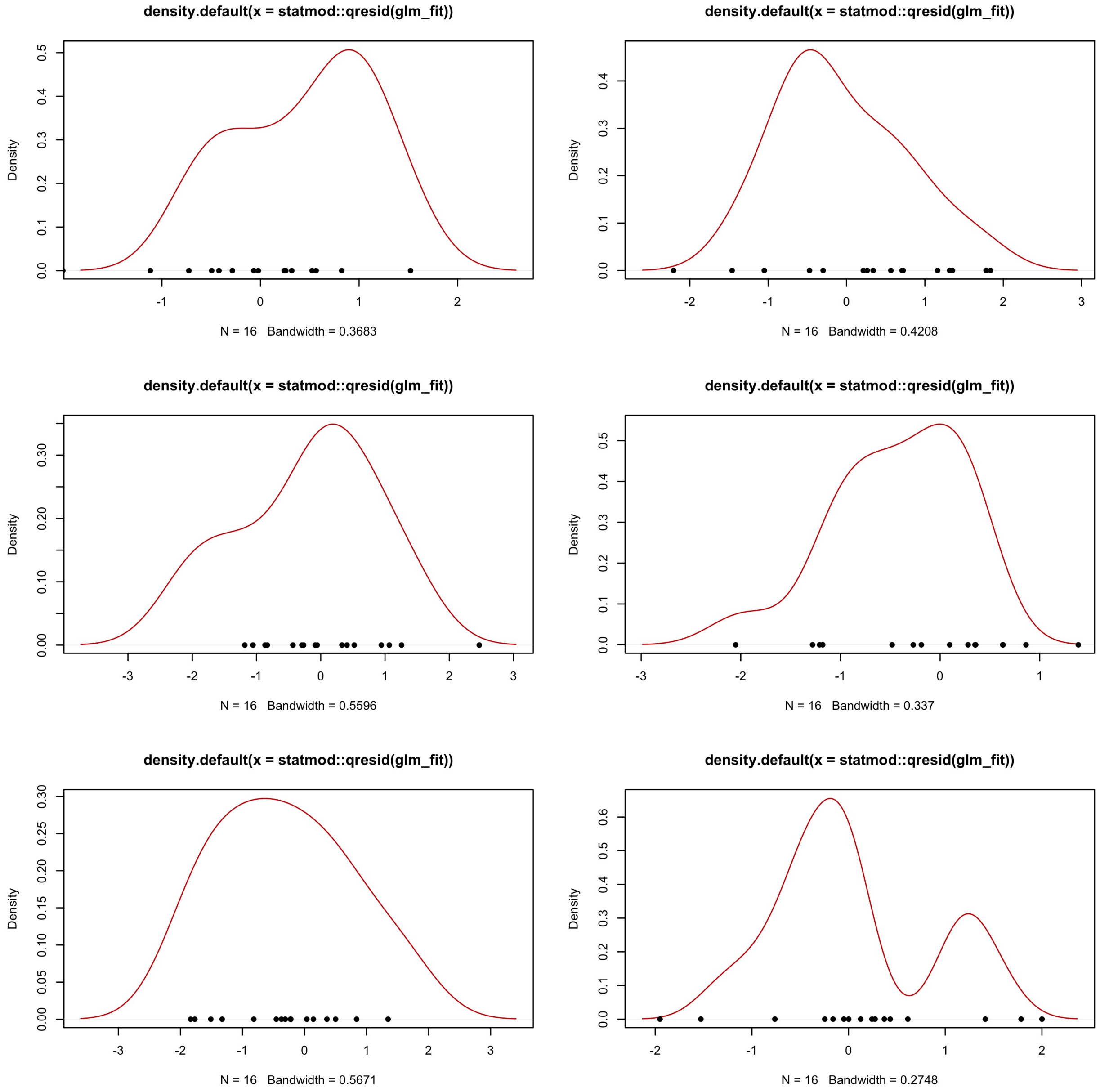


Figure M6: Density plots of 6 batches of randomized quantile residuals. Deviations from normality suggest violations of the exponential distribution assumption underlying logistic regression.

predictive modeling, we did not impute, and variables with missing (out-of-range) values were excluded. Third, we chose to exclude data obtained from an earlier batch used to test the assay. This batch included samples from two cases, one PGD+ and one PGD–. While it would have increased our sample sizes significantly, we were not prepared to incorporate batch effects into our analysis.

To test the robustness of our results to these choices, we re-ran the full analysis using three alternative settings, in each case holding other choices fixed: (1) using concentrations without imputations for exploratory analysis and with imputations for predictive modeling; (2) using fluorescence intensities (FIs) rather than calculated observed concentrations; and (3) including data for two additional cases assayed in a different batch.

We compared the results obtained in both robustness checks to those obtained in the main analysis to identify any qualitative differences, and we calculated the differences in optimal performance between corresponding model families and predictor sets.

Kolde, Raivo. 2019. "Pheatmap: Pretty Heatmaps."

Snell, Gregory I., Roger D. Yusen, David Weill, Martin Strueber, Edward Garrity, Anna Reed, Andres Pelaez, et al. 2017. "Report of the ISHLT Working Group on Primary Lung Graft Dysfunction, Part I: Definition and Grading—A 2016 Consensus Group Statement of the International Society for Heart and Lung Transplantation." *The Journal of Heart and Lung Transplantation* 36 (10): 1097–1103. https://doi.org/10.1016/j.healun.2017.07.021.

Wei, Taiyun, Viliam Simko, Michael Levy, Yihui Xie, Yan Jin, Jeff Zemla, Moritz Freidank, Jun Cai, and Tomas Protivinsky. 2021. "Corrplot: Visualization of a Correlation Matrix."
