## Supplemental Table 1 for "Predicting Primary Graft Dysfunction in Lung Transplantation: Machine Learning–Guided Biomarker Discovery"

**Supplemental Table 1:** Full list of analytes screened in 71-multiplex assay including analyte abbreviations and lower/upper limits of assay detection.

| Analyte | Abbreviation | Lower Limit of Detection (pg/mL) | Upper Limit of Detection (pg/mL) |
|---|---|---|---|
| Secondary lymphoid-tissue chemokine / CCL21 | **6CKine** | 4.88 | 1404.45 |
| B-cell attracting chemokine 1 / CXCL13 | **BCA-1** | 0.24 | 1000 |
| Cutaneous T cell attracting chemokine / CCL27 | **CTACK** | 1.22 | 274.24 |
| Epidermal growth factor | **EGF** | 0.64 | 123.48 |
| Epithelial neutrophil activating peptide 78 / CXCL5 | **ENA-78** | 4.88 | 2052.09 |
| Eosinophil chemotactic protein 1 / CCL11 | **Eotaxin** | 0.64 | 37.39 |
| Eosinophil chemotactic protein 2 / CCL24 | **Eotaxin-2** | 2.44 | 904.29 |
| Eosinophil chemotactic protein 3 / CCL26 | **Eotaxin-3** | 12.21 | 39.3 |
| Fibroblast growth factor 2 | **FGF-2** | 5.12 | 5222.21 |
| Fms-related tyrosine kinase 3 ligand | **FLT-3L** | 0.19 | 10.58 |
| Chemokine CX3C ligand 1 / CX3CL1 | **Fractalkine** | 6.4 | 1044.64 |
| Granulocyte colony stimulating factor / CSF3 | **G-CSF** | 0.96 | 13745.48 |
| Granulocyte macrophage colony stimulating factor / CSF2 | **GM-CSF** | 0.51 | 307.15 |
| Growth regulated alpha protein / CXCL1 | **GROα** | 0.26 | 3920.3 |
| T lymphocyte secreted protein I-309 / CCL1 | **I-309** | 0.49 | 4.64 |
| Interferon alpha 2 | **IFN-α2** | 1.6 | 652.33 |
| Interferon gamma | **IFNγ** | 0.26 | 237.38 |
| Interleukin 10 | **IL-10** | 0.51 | 756.55 |
| Interleukin 12 subunit p40 | **IL-12p40** | 1.28 | 717.02 |
| Interleukin 12 subunit p70 | **IL-12p70** | 0.64 | 152.61 |
| Interleukin 13 | **IL-13** | 1.28 | 3458.11 |
| Interleukin 15 | **IL-15** | 0.64 | 131.64 |
| Interleukin 16 | **IL-16** | 2.44 | 1067.09 |
| Interleukin 17A | **IL-17A** | 0.26 | 94.74 |
| Interleukin 17E / Interleukin 25 | **IL-17E/IL-25** | 8 | 36821.86 |
| Interleukin 17F | **IL-17F** | 6.4 | 129.43 |
| Interleukin 18 | **IL-18** | 0.13 | 7461.43 |
| Interleukin 1 receptor antagonist | **IL-1RA** | 0.32 | 555.66 |
| Interleukin 1 alpha | **IL-1α** | 0.96 | 543.73 |
| Interleukin 1 beta | **IL-1β** | 0.32 | 2108.53 |
| Interleukin 2 | **IL-2** | 0.13 | 37.3 |
| Interleukin 20 | **IL-20** | 12.21 | 232.13 |
| Interleukin 21 | **IL-21** | 4.88 | 13.93 |
| Interleukin 22 | **IL-22** | 2.56 | 887.71 |
| Interleukin 23 | **IL-23** | 12.21 | 1130.2 |
| Interleukin 27 | **IL-27** | 3.2 | 4732 |
| Interleukin 28A | **IL-28A** | 2.44 | 41 |

| | | | |
|---|---|---|---|
| Interleukin 3 | **IL-3** | 0.26 | 3 |
| Interleukin 33 | **IL-33** | 4.88 | 61 |
| Interleukin 4 | **IL-4** | 0.13 | 41 |
| Interleukin 5 | **IL-5** | 0.13 | 10 |
| Interleukin 6 | **IL-6** | 0.13 | 3968 |
| Interleukin 7 | **IL-7** | 0.13 | 63 |
| Interleukin 8 | **IL-8** | 0.13 | 9217 |
| Interleukin 9 | **IL-9** | 0.13 | 44 |
| Interferon gamma induced protein 10 / CXCL10 | **IP-10** | 0.51 | 9195 |
| Leukemia inhibitory factor / LIF interleukin 6 family cytokine | **LIF** | 4.88 | 10 |
| Monocyte chemoattractant protein 1 / CCL2 | **MCP-1** | 0.64 | 3499 |
| Monocyte chemoattractant protein 2 / CCL8 | **MCP-2** | 1.22 | 190 |
| Monocyte chemoattractant protein 3 / CCL7 | **MCP-3** | 1.6 | 231 |
| Monocyte chemoattractant protein 4 / CCL13 | **MCP-4** | 2.44 | 57 |
| Macrophage colony stimulating factor / CSF1 | **M-CSF** | 8 | 1252 |
| Macrophage derived chemokine / CCL22 | **MDC** | 0.13 | 1028 |
| Monokine induced by gamma interferon / CXCL9 | **MIG** | 1.28 | 49697 |
| Macrophage inflammatory protein 1 alpha / CCL3 | **MIP-1α** | 0.64 | 5196 |
| Macrophage inflammatory protein 1 beta / CCL4 | **MIP-1β** | 0.08 | 436 |
| Macrophage inflammatory protein 1 delta / CCL15 | **MIP-1δ** | 12.21 | 367 |
| Platelet-derived growth factor A | **PDGF-AA** | 2.56 | 765 |
| Platelet-derived growth factor B | **PDGF-AB/BB** | 1.92 | 36738 |
| Regulated upon activation, normally T cell expressed and secreted / CCL5 | **RANTES** | 0.26 | 1579 |
| Soluble CD40 ligand | **sCD40L** | 2.56 | 4924 |
| Stem cell factor / KIT ligand | **SCF** | 2.44 | 71 |
| Stromal cell derived factor 1 alpha and beta / CXCL12α+β | **SDF-1α+β** | 24.41 | 7183 |
| Thymus and activation regulated chemokine / CCL17 | **TARC** | 0.24 | 163 |
| Transforming growth factor alpha | **TGFα** | 0.26 | 74 |
| Tumor necrosis factor alpha | **TNFα** | 1.28 | 757 |
| Tumor necrosis factor beta | **TNFβ** | 0.32 | 363 |
| Thrombopoietin | **TPO** | 12.21 | 256 |
| TNF related apoptosis inducing ligand / CD253 | **TRAIL** | 2.44 | 980 |
| Thymic stromal lymphopoietin | **TSLP** | 2.44 | 6 |
| Vascular endothelial growth factor A | **VEGF-A** | 0.51 | 188 |
